# The ataxia protein sacsin is required for integrin trafficking and synaptic organization

**DOI:** 10.1101/2021.08.20.456807

**Authors:** Lisa E.L. Romano, Wen Yih Aw, Kathryn M. Hixson, Tatiana V. Novoselova, Tammy M. Havener, Stefanie Howell, Charlotte L Hall, Lei Xing, Josh Beri, Suran Nethisinghe, Laura Perna, Abubakar Hatimy, Ginevra Chioccioli Altadonna, Lee M. Graves, Laura E. Herring, Anthony J. Hickey, Konstantinos Thalassinos, J. Paul Chapple, Justin M. Wolter

## Abstract

Autosomal recessive spastic ataxia of Charlevoix-Saguenay (ARSACS) is caused by mutations in *SACS*, which manifest as a childhood-onset cerebellar ataxia. Cellular ARSACS phenotypes include mitochondrial dysfunction, intermediate filament (IF) disorganization, and loss of Purkinje neurons. It is unclear how the loss of SACS causes these deficits, or why they manifest as cerebellar ataxia. We employed a multi-omics approach to characterize molecular and cellular deficiencies in *SACS* knockout (KO) cells. We identified alterations in microtubule structure and dynamics, protein trafficking, and mislocalization of synaptic and focal adhesion proteins. Targeting *PTEN*, a negative regulator of focal adhesions, rescued several cellular phenotypes in *SACS* KO cells. We found sacsin interacts with proteins implicated in vesicle transport, including HSP proteins, and interactions between structural and cell adhesion proteins were diminished in *SACS* KO cells. In all, this study suggests that trafficking and localization of synaptic adhesion proteins is a causal molecular deficiency in ARSACS.

## Introduction

ARSACS is a childhood-onset neurological disease characterized by pyramidal spasticity, cerebellar ataxia, and Purkinje cell loss, that is thought to have both neurodegenerative and neurodevelopmental components^1^. Initially thought to be localized to the Charlevoix-Saguenay region of Quebec, Canada, over 170 distinct mutations have now been identified, making ARSACS the second most common recessive cerebellar ataxia worldwide^2^.

The large size of SACS (~520 kDa) has hampered biochemical and structural investigations into its function. SACS contains multiple domains that link it to molecular chaperone and protein quality control systems, including a ubiquitin like domain, three repetitive regions with some homology to the ATPase domain of Hsp90, and a J-protein domain, suggesting the potential to recruit Hsp70 chaperone proteins. Patient derived fibroblasts and the ARSACS mouse model possess consistent cellular phenotypes, including mitochondrial dysfunction, vimentin bundling, accumulation of non-phosphorylated neurofilament heavy, and altered firing properties of Purkinje neurons^3–9^. However, each of these deficits are found in a wide range of neurodegenerative diseases with diverse molecular causes^10,11^. Thus, the basic molecular function of sacsin, and why it’s loss manifests as a cerebellar ataxia, remains unclear.

## Results

### Comprehensive proteomics of *SACS* KO cells

To understand the molecular deficiencies in ARSACS, we deleted *SACS* using CRISPR/Cas9 in the SH-SY5Y human cell line (Extended Data Fig. 1a), which is widely used to model neurodegenerative diseases^12^. Consistent with ARSACS patient fibroblasts^4^ and *SACS*^(-/-)^ mice^7^, KO cells had abnormal bundling and asymmetric partitioning of multiple IFs, including vimentin (Fig. 1a), neurofilament heavy, and peripherin (Extended Data Fig. 1b-e). However, whether abnormal IF structure plays a causal role in ARSACS pathology is unclear. As phosphorylation is a key modification controlling IF assembly and disassembly^13^, we performed quantitative proteomic and phosphoproteomic profiling of *SACS* KO cells (Supplementary Table 1). We identified changes in several proteins previously described in ARSACS patient fibroblasts, including vimentin, the mitochondrial protein ATP5J, and the autophagy regulated scaffold SQSMT1/p62^4^ (Fig. 1b). Among the overabundant proteins were the tau-tubulin kinase 1 (TTBK1) and microtubule associated protein tau (MAPT) (Fig. 1c-f), which was hyperphosphorylated at several sites (Extended Data Fig. 1f, Fig. 1g,h, Supplementary Table 1). To assess the functional significance of specific phosphosites, we analyzed our data in light of a recent machine learning approach which estimated the effects of individual phosphosites on organism fitness^14^. This analysis identified several highly functional hypophosphorylated residues in vimentin, and the nuclear lamina IFs LMNA/B2, which is intriguing considering that ARSACS neurons have altered nuclear shape and positioning^4^ (Supplementary Table 1, Fig. 1i). Other hypophosphorylated proteins included the focal adhesion protein zyxin, and ataxin 2-like protein. In addition to tau, several other microtubule regulating proteins were hyperphosphorylated, including the primary cilia protein ARL3^15^, and the scaffold stathmin, which promotes microtubule assembly in a pS16 dependent fashion^16^.

**Figure 1.**
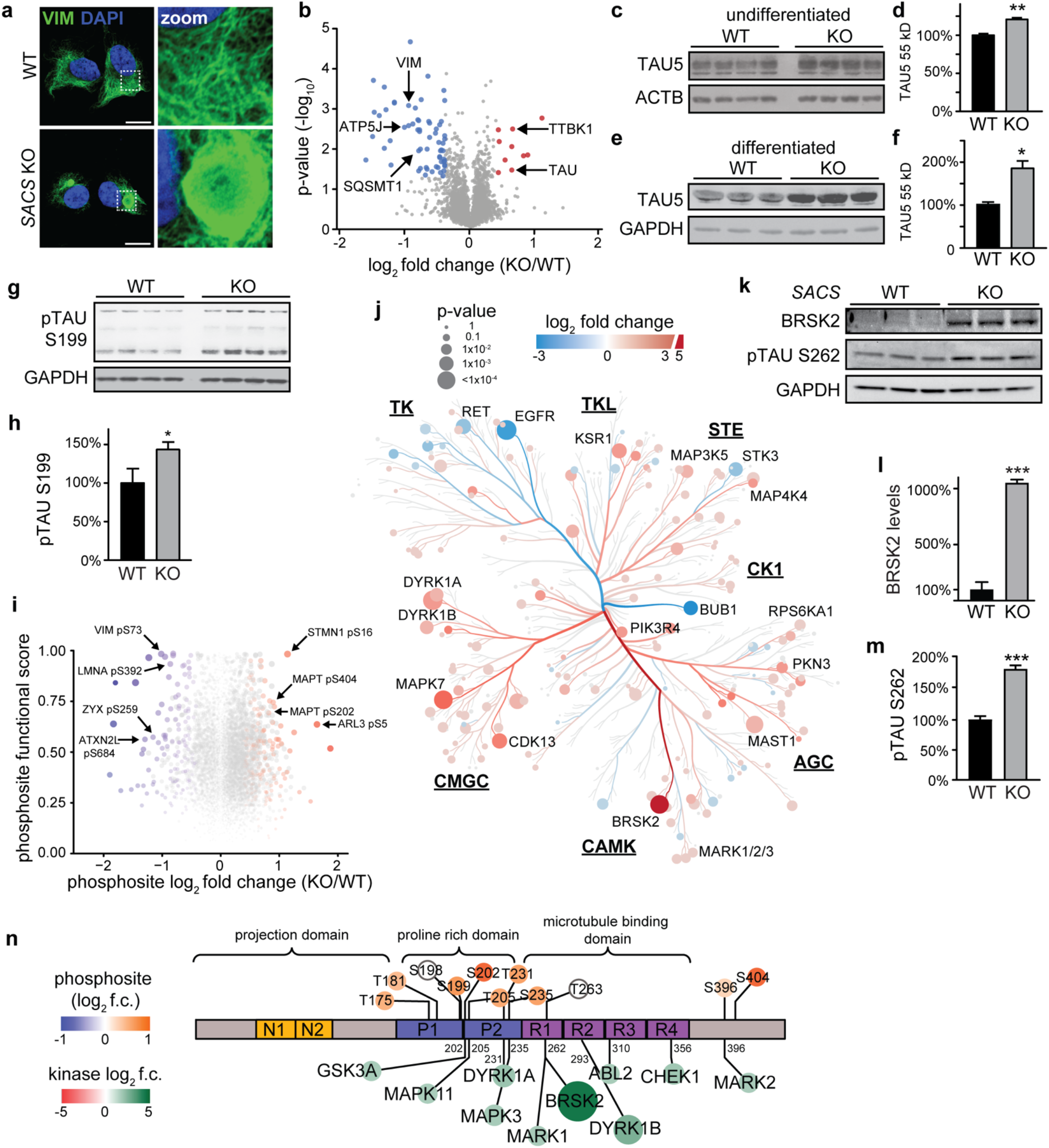
Proteomic profiling of *SACS* KO cells. a. Representative confocal images of control (WT) and *SACS* knockout (KO) SH-SY5Y neuroblastoma cells immunostained for the intermediate filament protein vimentin. Scale bar = 10 μm. b. Global proteomic profiling of *SACS* KO SH-SY5Y cells (*SACS* KO). Cutoffs for significance were p<0.05 and log_2_ fold change -/+0.4. c-f. Western Blot analysis quantification of pan-tau (Tau5) in *SACS* KO and WT cells in undifferentiated (c,d) and neuronally differentiated (e,f) SH-SY5Y cells. g,h. Western blot and quantification of phosphorylated tau at serine 199.Phosphoproteomic analysis of *SACS* KO SH-SY5Y cells. Green circles mark specific phosphorylated residues on tau. i. Functional analysis of altered phosphosites in *SACS* KO cells. Y-axis is the functional score assigned by Ochoa et al (2020), which combines 59 features to assess the imact of each phosphosite on organismal fitness. Dot color and size reflects a combined metric of log_2_ fold change and functional score (product of the two together). Non-significant phosphosites are colored grey. j. Phylogenetic tree of the kinome in *SACS* KO cells. 35% of kinases were altered in *SACS* KO cells (log_2_ f.c. -/+0.4, p<0.05). Color indicates log_2_ f.c. of kinase abundance, size indicates -log_10_ p-value. Underlined abbreviations refer to phylogenetically related kinase families. k-m. Western blot and quantification for BRSK2, and the BRSK2 target residue pTAU Ser262. n. Protein map of tau isoform 2 (2N4R). Phosphosites identified in phosphoproteomic profiling are labelled above diagram. Tau kinases identified in the kinome profiling are listed below, labeled with known amino acid substrates in tau. Colored circles correlate with log_2_FC of significantly differentially expressed phosphosites or kinases. Unless otherwise noted, all error bars are S.E.M., all statistical tests are Student’s t-test (*P<0.05, **P<0.01, ***P<0.001, ****P<0.0001).

Kinases are attractive drug targets^17^, but are typically lowly expressed and difficult to detect with standard proteomics. Therefore, we enriched for kinases using multiplexed kinase inhibitor beads, and performed quantitative mass-spectrometry^18^. The kinome was broadly altered in *SACS* KO cells (Extended Data Fig. 1g,h, Supplementary Table 1), with affected kinases spread among all kinase families (Fig. 1j). Interestingly, specific families were generally misexpressed in similar direction, for example the tyrosine kinase family (TK) members were generally downregulated, while CMGC family members were generally upregulated (Fig. 1j). The most overabundant kinase, BRSK2, and additional CAMK family members MARK1/2/3, all phosphorylate Ser262 in the microtubule binding domain of tau^19,20^ (Fig. 1k-m). Thr231, which is phosphorylated by DYRK1A, is also associated with the detachment of tau from microtubules^21,22^. In all, there were 10 overexpressed kinases which phosphorylate tau at specific sites (Fig. 1n). In pathological settings, tau overabundance and hyperphosphorylation causes elevated microtubule stability, interferes with motor protein function, and alters axonal trafficking^23–25^. Combined with the altered phosphorylation of other microtubule related proteins (Fig. 1i), this data suggests that microtubule structure or function may be altered in ARSACS.

### Microtubule organization and dynamics are altered in *SACS* KO cells

There is a bidirectional relationship between vimentin and microtubules, where vimentin filaments form along microtubule tracts, and can facilitate microtubule nucleation^26^. We found that cage-like vimentin bundles form around gamma-tubulin, a marker of the microtubule organizing center (MTOC), which is a central hub for microtubule nucleation and cargo transport^27^ (Fig. 2a, Extended Data Fig. 2a). Acetylated alpha-tubulin, a microtubule stabilizing post-translational modification, was increased in *SACS* KO cells, without affecting total alphatubulin distribution or levels (Extended Data Fig. 2b, Fig. 2b,c). To assess microtubule dynamics we treated cells with the microtubule destabilizer nocodazole, and found enhanced microtubule polymerization following nocodazole washout (Fig. 2d,e). *SACS* KO cells also demonstrated increased microtubule polymerization and disordered movements as assessed by live cell imaging of the microtubule plus-end binding protein EB1:GFP (Fig. 2f, Extended Data Fig. 2e, Supplementary Video 1,2).

**Figure 2.**
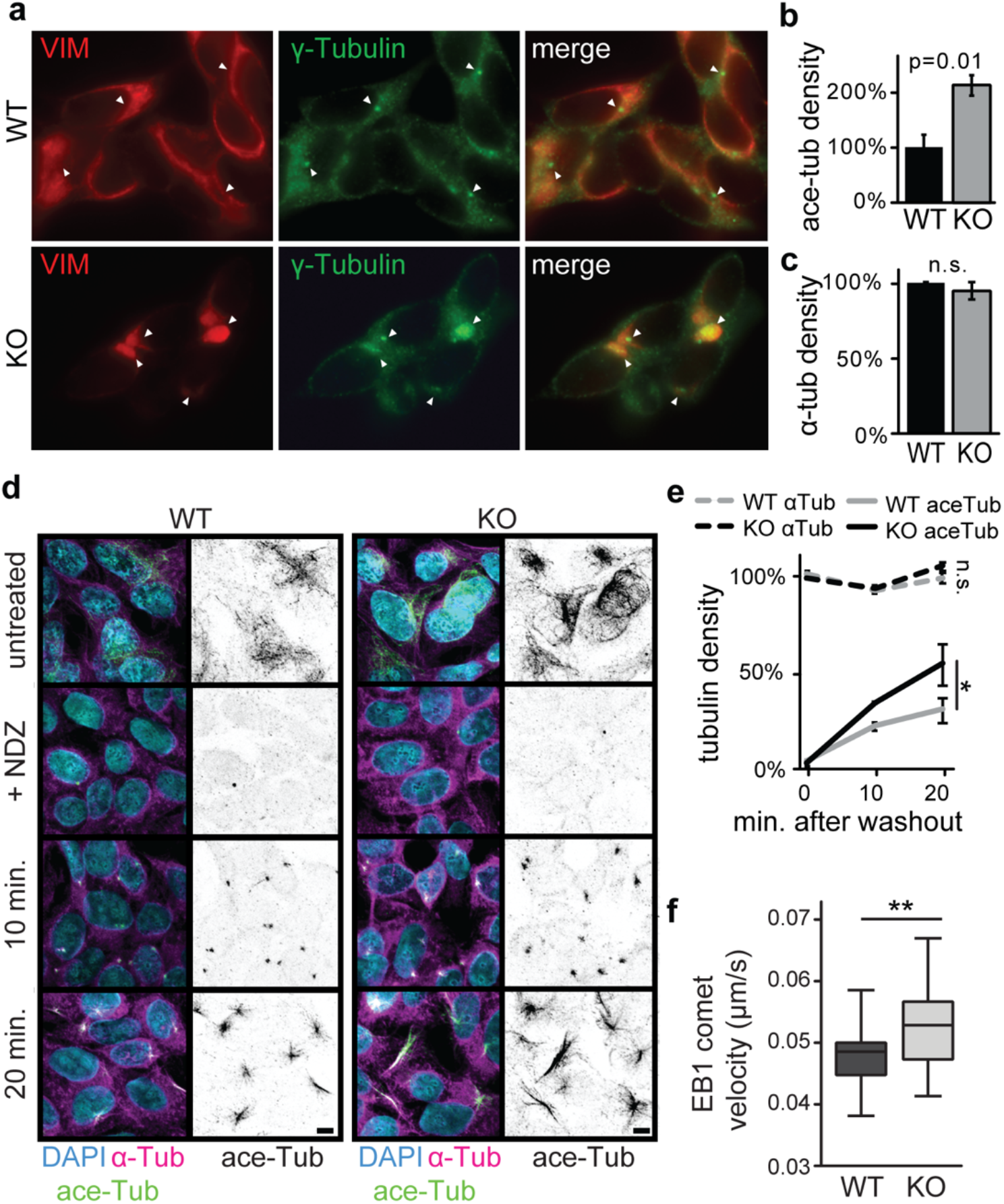
Altered microtubule structure and dynamics in *SACS* KO cells. a. Confocal immunofluorescent images of *SACS* WT/KO cells stained for vimentin, and the MTOC marker gamma tubulin. Arrowheads point to the most intense signal in each cell, showing that vimentin bundles surround the MTOC in *SACS* KO cells. Scale bar = 10 μm. b-c. Quantification of images in Extended Data Fig. 2b, showing altered microtubule acetylation in *SACS* KO cells (b) without global changes in microtubule structure (c). d. Confocal images of WT/KO cells treated with nocodazole (NDZ) labeled for alpha- and acetylated-tubulin at indicated time points following nocodazole washout. Note the faster microtubule repolymerization and acetylation in *SACS* KO cells. Scale bar = 10 μm. e. Quantification of images in (d). n = 3 coverslips; One-way ANOVA with Tukey post-test. f. Quantification of microtubule polymerization velocity marked by EB1-GFP movement in WT/KO cells on TIRF microscope from Supplemental Video 1. n=34 WT and n=25 *SACS* KO cells, examined from at least three independent experiments; unpaired t-test.

Mitochondrial movement in neurons is dependent on microtubules^28^. Tau overexpression and hyperphosphorylation can cause decreased mitochondria trafficking along microtubules^19,29,30^, build-up of mitochondria around the MTOC^31^, and DRP1 mislocalization and reduced mitochondrial fission^32,33^. In ARSACS, mitochondria also accumulate around proximal dendrites^6^ and exhibit reduced DRP1 dependent fission^3^. We observed occlusion of mitochondria around vimentin bundles (Extended Data Fig. 2c), with no alterations in the actin cytoskeleton^4^ (Extended Data Fig. 2d). To assess how these alterations affect mitochondria in neurons, we performed neuronal differentiation of SH-SY5Y cells^34^. WT and *SACS* KO cells expressed indistinguishable levels of neuronal markers, suggesting equal differentiation capacity (Extended Data Fig. 2f,g), but neurites were fewer and shorter in *SACS* KO cells (Extended Data Fig. 2f-i). *SACS* KO neurites also possessed fewer mitochondria (Extended Data Fig. 2j), and diminished mitochondrial movement (Extended Data Fig. 2k, Supplementary Video 3). Our proteomics data also identified hyperphosphorylation in several kinesin proteins, including multiple residues on KIF1A (Supplementary Table 1), which shuttle mitochondria along microtubule tracts^35^. In all, these results demonstrate altered microtubule structure, dynamics, and function in *SACS* KO cells, and are consistent with the microtubule stabilizing effects of tau and STMN1 pS16 hyperphosphorylation^16^.

### Focal adhesion organization and dynamics are disrupted in *SACS* KO cells

To more systematically characterize our proteomic datasets, we performed gene ontology (GO) analysis (Fig. 3a,b, Supplementary Table 2). The top associated terms were related to ‘focal adhesions’, including ‘integrin signaling’, ‘cell-matrix adhesions’, and ‘cadherin binding’. Focal adhesions are plasma membrane-associated macromolecular assemblies that physically link the intracellular cytoskeleton and extracellular matrix (ECM), and function as signaling nodes to regulate force dependent cellular processes^36^. Focal adhesions are composed of integrin receptors bridging the ECM with actin bundles, which interact with microtubules and IFs to coordinate dynamic regulation of focal adhesion structure^37–39^. In the brain, focal adhesions are critical for structural remodeling during axon growth and synapse formation and maintenance^40^.

**Figure 3.**
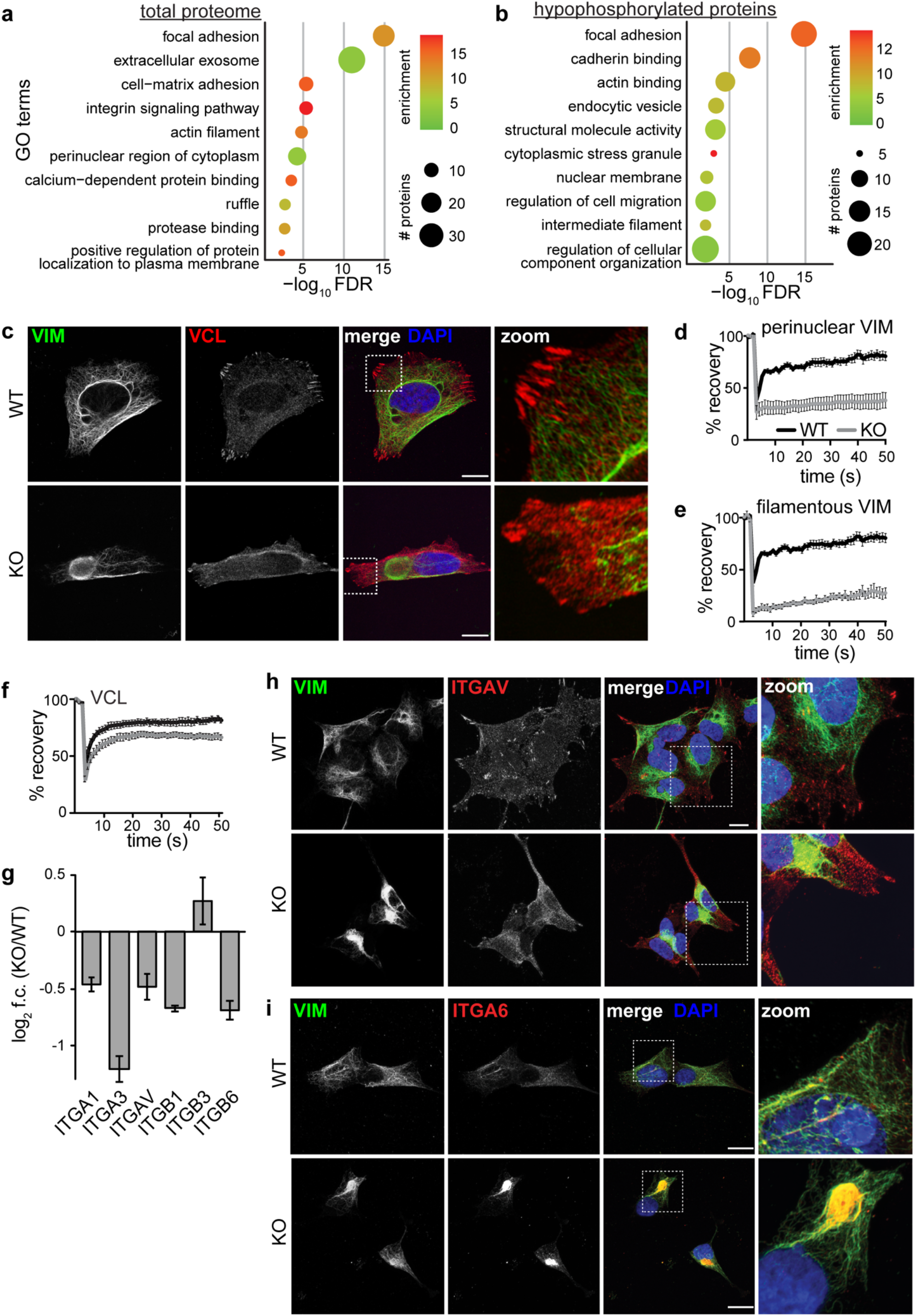
Focal Adhesions are altered in *SACS* KO cells. a,b. GO term analysis from the total proteome (p<0.05, log_2_ fold change cutoff -/+0.4) (a), and hypophosphorylated (p<0.05, log_2_ fold change <-0.4) ( (b) proteins. Both identify focal adhesions as the main process that is altered in *SACS* KO cells. c. Confocal images of WT/KO SH-SY5Y cells immunolabeled for vimentin and the focal adhesion protein vinculin. d-f. FRAP analysis of perinuclear vimentin (d), filamentous vimentin on the periphery of the cell away from vimentin bundle (e), and the focal adhesion protein vinculin (f). Cells were transfected with tomato-VIM/VCL expression vector and defined 2 × 2 μm regions of interest were bleached by using a 568-nm laser line. Recovery was monitored over 50 cycles of imaging with a 1-s interval. n=10 cells from each of three independent experiments. g. Changes in levels of integrin proteins quantified by mass-spectrometry (Fig. 1b). h,i. Representative confocal images of cells immunolabeled for ITGAV (h) and ITGA6 (i). Scale bar = 10 μm.

We observed decreased focal adhesion number, area, and aspect ratio in *SACS* KO cells by immunolabelling for core focal adhesion components paxillin and vinculin, whose total levels were unaffected (Fig. 3c, Extended Data Fig. 3a-h, Supplementary Table 1). While paxillin is primarily localized at focal adhesions, it also is known to interact with the MTOC^41^, and we observed perinuclear accumulation of paxillin coinciding with the vimentin bundle (Extended Data Fig. 3a). Microtubules regulate vinculin localization to focal adhesions^42^, and we found reduced vinculin and vimentin dynamics in *SACS* KO cells using fluorescence recovery after photobleaching (FRAP) (Fig. 3d-f). We next removed cell bodies with hypotonic shock, leaving only the structural remnants of cell/ECM interactions, and again found reduced vinculin structures, suggesting that the mislocalization of adhesion proteins results in decreased cell/ECM interactions (Extended Data Fig. 3i-l). These findings were consistent in *SACS* KO HEK293 cells, which were generated using an alternate CRISPR/Cas9 genome editing strategy^4^ (Extended Data Fig. 3m-q). Our proteomics data also revealed changes in several integrin proteins (Fig. 3g). Localization of ITGAV to focal adhesions was lost in *SACS* KO cells (Fig. 3h), while ITGA6 was sequestered in the vimentin bundle (Fig. 3i).

### Modulating PTEN-FAK signaling rescues cellular deficits in *SACS* KO cells

Beyond providing structural support for cells, focal adhesions are enriched with many signaling proteins, which transmit signals from the extracellular milieu to effectors in the cytoplasm and nucleus. A master regulator of focal adhesion signaling is the focal adhesion kinase (FAK/PTK2)^43^, which regulates neuronal outgrowth and synapse formation by phosphorylating downstream effectors of focal adhesion signaling^44^ (Fig. 4a). Although total levels of FAK were unaltered, phosphorylated FAK (pFAK) was significantly reduced in *SACS* KO cells (Fig. 4b, Extended Data Fig. 4a,b). Similarly, JNK and paxillin were also hypophosphorylated, while total levels of these proteins were unaffected (Fig. 4b, Extended Data Fig. 4c-g, Supplementary Table 1). Localization of pFAK to focal adhesions was also decreased *SACS* KO cells (Fig. 4c). We next considered the mechanism by which FAK signaling is suppressed in *SACS* KO cells. The phosphatase PTEN, which dephosphorylates FAK and negatively regulates FAK activity^45^, was elevated in *SACS* KO cells (Fig. 4b, Extended Data Fig. 4h). To investigate whether increased PTEN is a general consequence of IF disorganization, we treated WT SH-SY5Y cells with simvastin^46^, which induced bundling and perinuclear accumulation of vimentin, but did not affect PTEN levels (Extended Data Fig. 4i-k). Conversely, reducing *PTEN* to WT levels in *SACS* KO cells (Extended Data Fig. 4l,m) increased pFAK (Fig. 4d), reduced the frequency of perinuclear vimentin accumulation (Fig. 4e,f), increased the number of focal adhesions (Fig. 4e,g), and restored migration deficits of *SACS* KO cells (Extended Data Fig. 4n,o). Together these results indicate that increased PTEN activity contributes, at least in part, to the IF and focal adhesion phenotypes in *SACS* KO cells. Furthermore, our data suggests that modulating this pathway may reduce molecular deficits associated with ARSACS.

**Figure 4.**
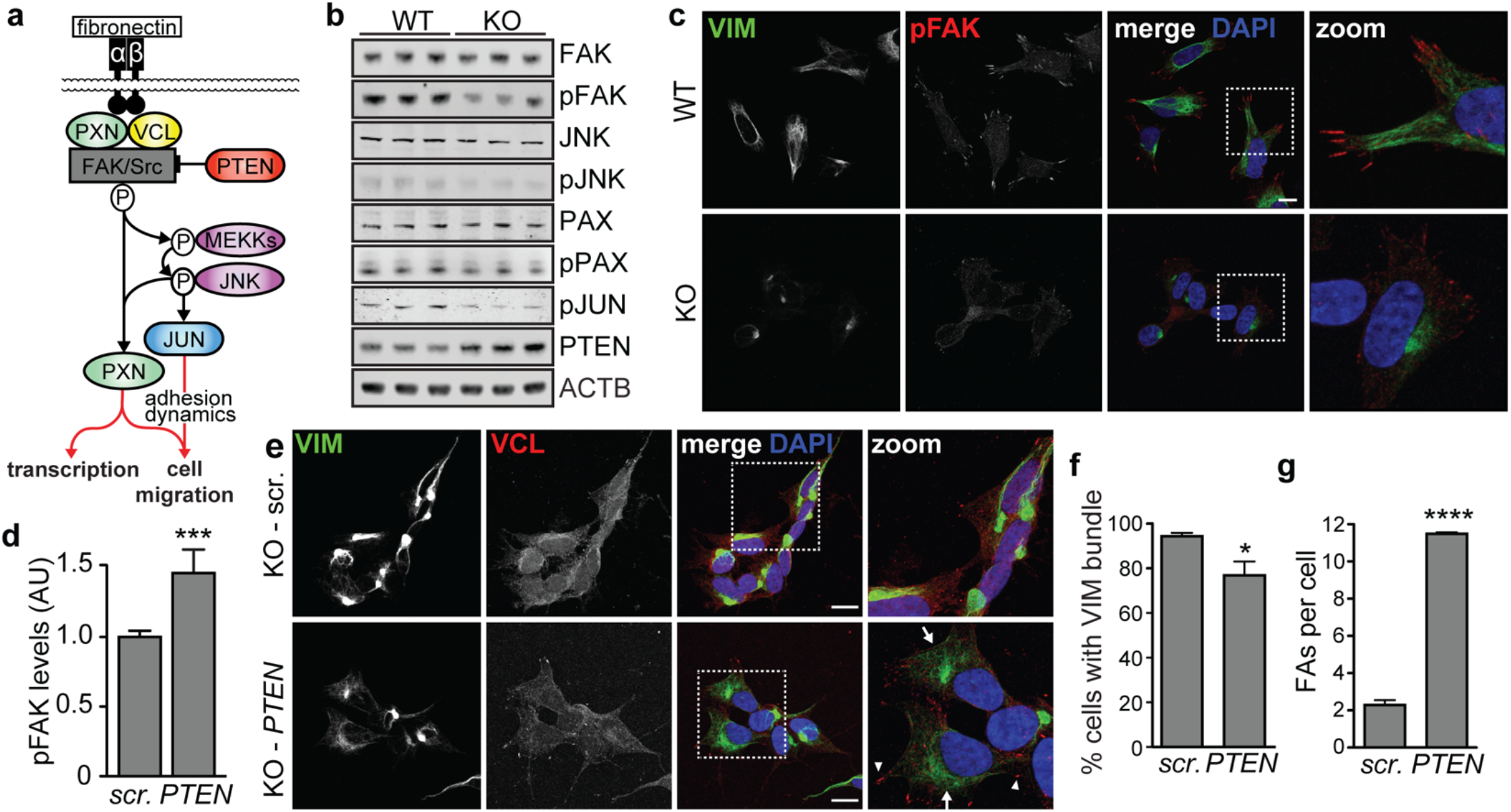
Targeting upstream focal adhesion regulator PTEN rescues focal adhesion and vimentin bundling phenotypes in *SACS* KO cells. a. Model of regulators and effectors of focal adhesion signaling b. Western blots for regulators PTEN, FAK, phosphorylated FAK (pFAK), JNK, phosphorylated JNK (pJNK), paxillin (PAX), phosphorylated paxillin (pPAX) and phosphorylated Jun (pJUN) in total cell lysates from *SACS* KO and control cells. β-actin used to confirm equivalent sample loading. c. Representative confocal images of cells immunolabeled for pFAK. Scale bar = 10 μm. d. pFAK levels with *SACS* KO cells treated with either scrambled (scr.) or siRNA targeting *PTEN*. e. Representative confocal images for cells transfected with siRNAs targeting PTEN (*PTEN*) or scramble siRNAs and immunolabeled for vimentin and vinculin. Arrowheads in the zoomed panel indicate cells with prominent focal adhesions, arrows indicate cells with absent or reduced perinuclear accumulations of vimentin. Scale bars = 10 μm. f,g. Quantification of the incidence of sacsin KO cells with perinuclear accumulations of vimentin (f) or vinculin positive focal adhesions (g) 48 hours after transfection with siRNAs targeting *PTEN* or scr. siRNAs. n=3 replicates with >100 cells in each replicate.

### Membrane bound synaptic adhesion molecules are mislocalized in *SACS* KO cells

Focal adhesions act as signal transduction hubs to integrate information from the outside of the cell to the inside. Some focal adhesion proteins, including paxillin and zyxin (Fig. 1i), can shuttle to the nucleus and function as transcriptional coregulators in a phosphorylation dependent manner^47,48^. GO term analysis of hyperphosphorylated proteins identified several terms related to transcriptional processes, including nuclear matrix, RNA binding, and transcriptional coregulator activity (Extended Data Fig. 5a, Supplementary Table 2). Therefore, we performed RNA-seq in neuronally differentiated SH-SY5Y cells to determine how structural changes affect the neuronal transcriptome (Extended Data Fig. 5b, Supplementary Table 3). We found 876 differentially expressed genes (FDR<0.05, f.c. -/+ 0.4), suggesting the loss of sacsin has profound effects on the transcriptome (Extended Data Fig. 5b). GO term analysis identified multiple terms related to membrane proteins and neuronal synapses, including ‘post-synaptic membrane’ and ‘axon terminus’ (Extended Data Fig. 5c). Protein interaction mapping^49^ revealed altered expression of multiple ECM proteins, integrins, and regulators of integrin activation (Extended Data Fig. 5d), suggesting that membrane dependent signaling cascades affect the transcription of functionally related genes.

Cell surface proteins are frequently underrepresented in proteomics experiments due to low expression and biochemical properties^50^. While 26% of all genes identified by RNAseq were detected by proteomics experiments, only 11 % of differentially expressed genes (DEGs) were detected by proteomics (Extended Data Fig. 5e), which is in agreement with GO term analyses suggesting alterations in membrane dependent processes. Therefore to enrich for surface proteins, we incubated live cells with biotin, which labels cellular and exosomal membrane/surface proteins, followed by neutravidin purification and analysis by quantitative mass-spectrometry^51^ (Fig. 5a, Supplementary Table 1). This approach identified an additional 870 proteins not in our proteomic dataset (Extended Data Fig. 5f, Fig. 5b). Proteins altered KO cells included several signaling receptors (FGFR1, LRP4, NRP2), and GTP binding proteins involved in signal transduction (GNG2, GNG8). The most overabundant membrane protein was neurofascin (NFASC), a neuronal adhesion protein that has been linked to movement disorders and cerebellar ataxia^52,53^ (Fig. 5c). We next compared membrane proteins found in both proteomic and surfaceome datasets, reasoning that conflicting levels between the surface and total protein levels could reflect improper membrane recycling, precocious membrane localization, or deficits in membrane-bound trafficking. Many proteins with altered surface levels showed no, or even opposing change in total protein levels (Fig. 5d). Among the most mislocalized proteins were synaptic adhesion proteins, including multiple integrins (ITGA1, ITGB1, ITGA3), neuronal cell adhesion molecules (L1CAM, NRCAM, CNTN1, LSAMP), the focal adhesion regulator RET/GFRA3 heterodimer, the microtubule binding protein DCX, and AHNAK, a 700 kDa scaffolding protein with diverse yet poorly understood function^54^ (Fig. 5d).

**Figure 5.**
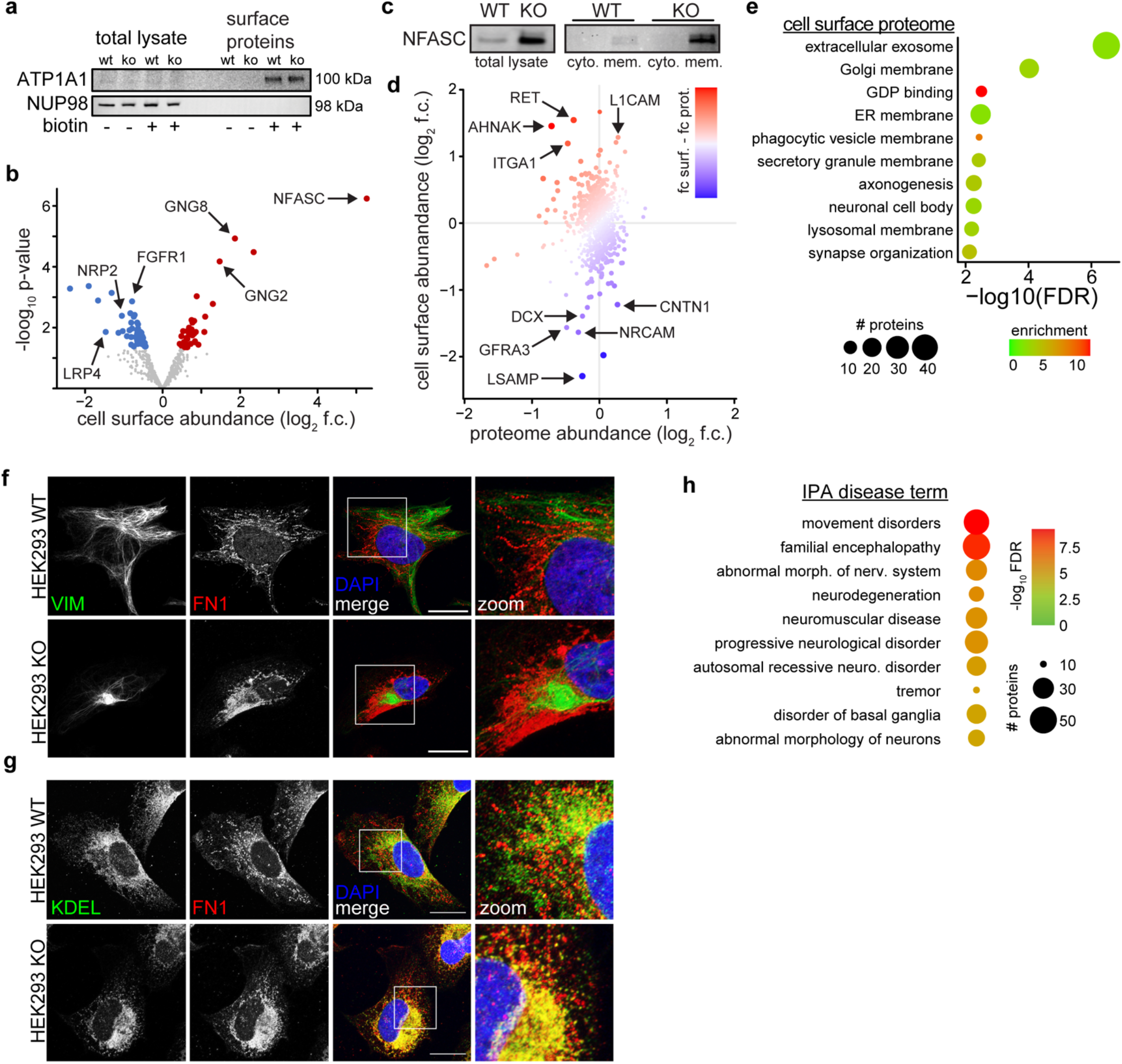
Cell surface proteomics suggests deficits in neuronal adhesion molecules is a causal molecular deficit in ARSACS. a. Western blot of membrane purification approach, illustrated by ATP1A1, a membrane bound Na/K ATPase, and NUP98, a nuclear pore protein. In total lysate only NUP98 is detectable. After purification ATP1A1 is detectable only in conditions that were treated with biotin, and NUP98 is not longer detected, suggested labelling specificity and enrichment of cell surface proteins. b. Volcano plot of cell surface proteins only detected in surface proteomic experiment. c. Western blot of NFASC in total lysate (left), and fractionated cytoplasmic or membrane fractions in WT and *SACS* KO cells. d. Levels of proteins detected in both cell surface and proteomic datasets. Proteins are colored by the disparity between these two datasets (fold change surface – fold chance proteome), with red indicating more, and blue less membrane abundance relative to total protein levels. e. GO term analysis of proteins differentially localized in membrane of *SACS* KO cells (p<0.05, log_2_ fold change -/+ 0.4). f,g. Representative confocal images for fibronectin (levels not affected in any proteomic experiment) and vimentin (f) and ER marker KDEL (g) in WT and *SACS* KO HEK293 cells. Scale bar = 10 μm. h. Disease enrichment analysis with Ingenuity Pathway Analysis (IPA) of significantly differentially expressed cell-surface proteins (p<0.05, log_2_ fold change -/+ 0.4).

GO term analysis of proteins with altered surface levels suggested deficits in processes related to vesicle packaging and transport (Fig. 5e). These included eight exosomal Rab proteins, which were increased in the surfaceome and not affected at the total protein level (Extended Data Fig. 5g, Supplementary Table 1). Rabs are a diverse family of GTPases that coordinate multiple aspects of membrane protein trafficking, including focal adhesion turnover, and integrin endo/exocytosis^55^. Specific Rabs also regulate trafficking between the Golgi and the endosomal network (RAB8A, RAB10), bidirectional Golgi/ endoplasmic reticulum (ER) trafficking (RAB2A, RAB18), and EGFR internalization (RAB7A)^56,57^. Kinome profiling also identified multiple regulators of Rab activity and trafficking, including PIK3R4 and PIK3C3, which regulate PTEN activity through localization to vesicles in a microtubule dependent fashion^58^ (Fig. 1j, Supplementary Table 1).

To assess trafficking and localization deficits independent of changes in protein level or phosphorylation, we investigated the localization of the ECM protein fibronectin, which is processed and packaged into vesicles in the ER and Golgi^59^, and trafficked to the cell periphery along microtubules^60^. In WT HEK293 cells fibronectin puncta were organized in ‘chains’, which appear collapsed around the vimentin bundle in *SACS* KO cells (Fig. 5f). Staining for the ER marker KDEL suggested that fibronectin is retained in the ER in HEK293 and SH-SY5Y *SACS* KO cells (Fig. 5g, Extended Data Fig. 5h), suggesting that membrane bound trafficking is substantially affected in *SACS* KO cells.

We next used Ingenuity Pathway Analysis (IPA) to assess whether the misregulated cell surface proteins are associated with any pathological conditions. Resoundingly, the terms were associated with disease traits reminiscent of ARSACS, including movement disorders, neurodegeneration, and progressive neurological disorder (Fig. 5h). Notably, three of the most mislocalized proteins, NFASC, NRCAM, and CNTN1, form molecular complexes that are important for axon guidance^61^, maintenance of synapses by astrocytes^62^, and interactions between Purkinje neuron axons and glia^63^. KO mice or humans which harbor mutations in each of these genes develop cerebellar ataxias with features that resemble ARSACS (see Discussion).

### Cell adhesion proteins are mislocalized in ARSACS mice

We next looked for evidence of changes in protein localization in *SACS^(-/-)^* mice at P120, which is during the initial period of Purkinje neuron loss^7^ (Fig. 6a). ITGA1, which was among the most mislocalized proteins in *SACS* KO cells (Fig. 5d), is normally localized along Purkinje axon tracts in unaffected mice, but in *SACS^(-/-)^* mice ITGA1 accumulated in the soma, dendritic trunk, and axonal swellings (Extended Data Fig. 6a, Fig. 6b). NFASC is normally enriched along the axon initial segment of Purkinje neurons where it contributes to action potential initiation^64^. *Sacs*^(-/-)^ Purkinje neurons showed increased NFASC intensity, as well as expanded localization around the circumference of the soma (Fig. 6c, Extended Data Fig. 6b). Purkinje neurons synapse onto neurons in the deep cerebellar nucleus (DCN), which is the primary output hub of the cerebellum. We observed striking loss of Purkinje neurons synapses onto DCN neurons in *Sacs*^(-/-)^ mice (Fig. 6d), which may be present at presymptomatic stages^9^. Colocalization between the presynaptic marker SYN1 and paxillin was only evident on DCN neurons in unaffected mice, but showed large extracellular accumulations in *SACS*^(-/-)^ mice (Extended Data Fig. 6b). We also observed abnormal axon termini from layer V cortical motor projection neurons in the spinal cord (Extended Data Fig. 7a,b), as well as smaller axonal bundles in *SACS^(-/-)^* mice (Extended Data Fig. 7c).

**Figure 6.**
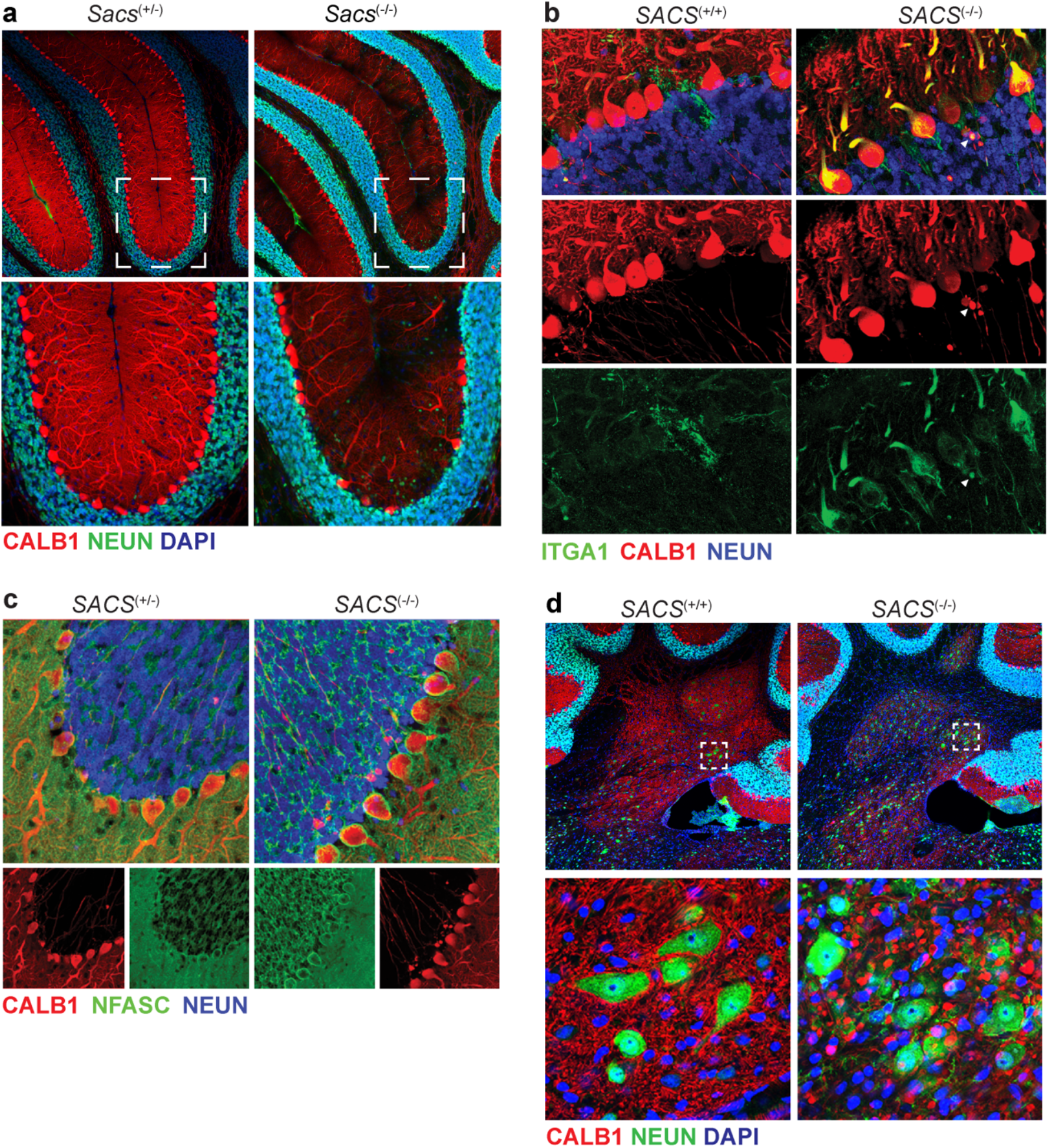
Altered localization of membrane proteins and synapses in ARSACS mice. a. Confocal imaging of Purkinje neurons in litter mate controlled P120 *SACS^(+/-)^* and *SACS^(-/-)^* mice. Purkinje marker calbindin (CALB1), neuronal marker (NEUN), and nuclei (DAPI). *SACS^(+/-)^* mice are phenotypically normal, analogous to unaffected human carriers. b. High magnification of Purkinje neuron layer in P120 mice, stained for integrin A1 (ITGA1), one of the most significantly mislocalized proteins in mass-spec (Fig. 6b,c). ITGA1, normally localized along axon tracts (Extended Data Fig. 6a), accumulated in initial dendritic segment in *SACS* KO mice. Arrowhead marks ITGA1 accumulation in axonal swellings, typical of ARSACS mice. c. High magnification of Purkinje neuron layer in P120 mice, stained for neurofascin (NFASC). NFASC is normally localized to the axon initial segment (AIS). In *SACS* KO mice NFASC staining is more intense around Purkinje soma, and more prominent around the entire soma, as opposed to primarily adjacent to the AIS. d. Confocal imaging of the DCN in P120 *SACS* KO mice, demonstrating synaptic deficits between Purkinje neurons and their synaptic termini on DCN neurons.

### The loss of sacsin disrupts protein-protein interactions

To identify how the loss of *SACS* causes abnormal protein trafficking, we performed quantitative label-free mass spectrometry of endogenous sacsin co-immunoprecipitated (co-IP) from WT SH-SY5Y cells. KO cells were also used to control for non-specific protein pulldown. Our analysis identified 90 proteins as putative Sacsin interactors, including vimentin and vinculin (Supplementary Table 4). Immunofluorescence revealed sacsin puncta in and around vinculin positive focal adhesions (Extended Data Fig. 8a,b), and in close proximity to vimentin structures, with sacsin often being between them (Fig. 7a). Reciprocal co-IP experiments confirmed interactions between sacsin, vimentin and vinculin, but the interaction between vimentin and vinculin was dramatically reduced in *SACS* KO cells (Fig. 7b). NFASC has been reported to interact with vimentin^65^, leading us to wonder whether NFASC may also interact with focal adhesion proteins. Co-IP experiments identified an interaction between NFASC and vinculin, which was dramatically reduced in *SACS* KO cells (Fig. 7c), suggesting that sacsin promotes the formation and/or stabilization of adhesion protein interactions.

**Figure 7.**
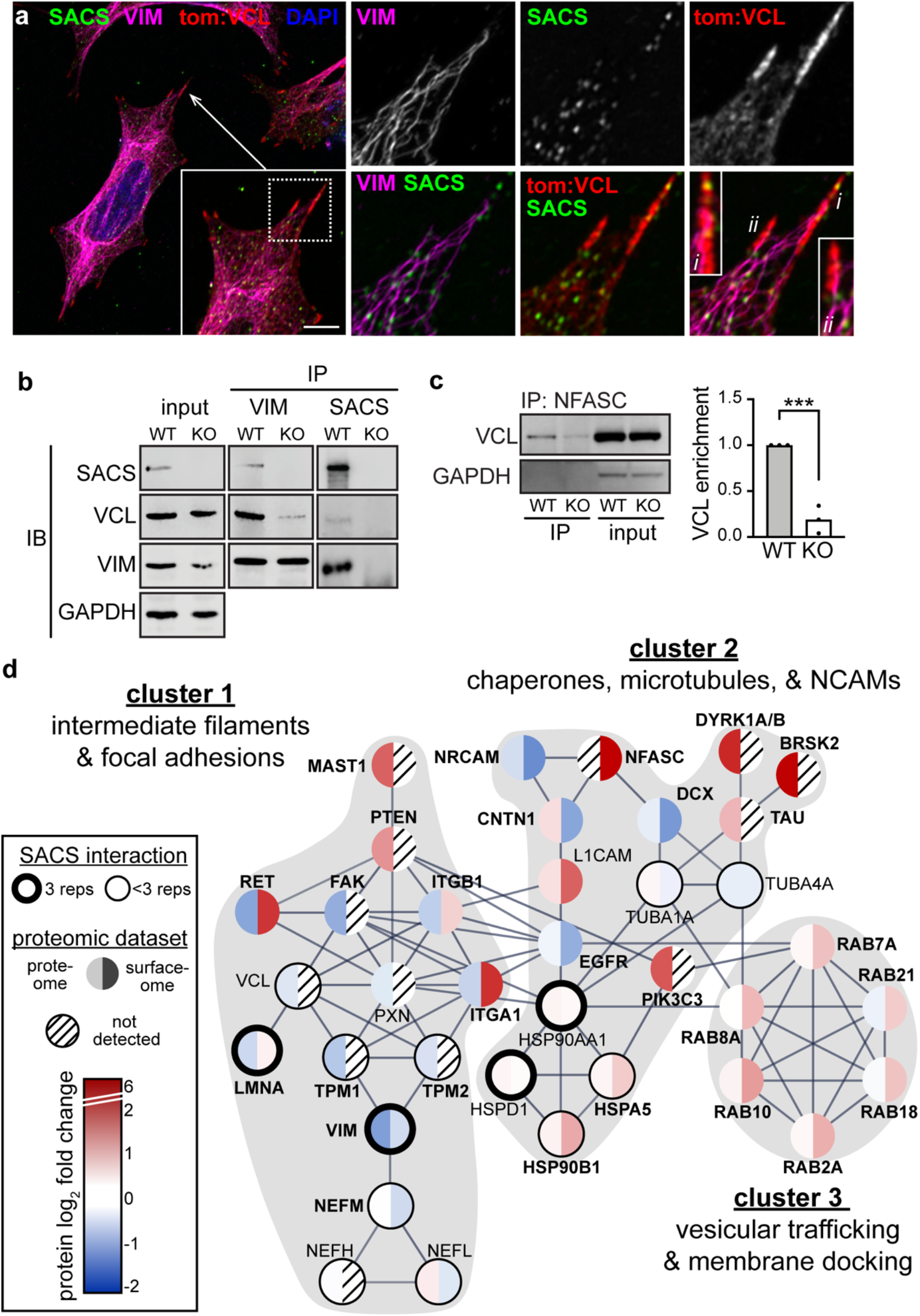
The loss of Sacsin disrupts protein-protein interactions. a. Representative Airyscan confocal analysis of sacsin, vimentin, and transfected tdTomato:vinculin staining in WT SH-SY5Y cells, demonstrating SACS localization along vimentin tracts and focal adhesions. Scale bar = 10 μm. b. Vimentin or sacsin were immunoprecipitated from WT and *SACS* KO SH-SY5Y cells, and coimmunoprecipitated proteins (sacsin, vinculin, vimentin) were analyzed by Western blot. Results suggest decreased interaction between vimentin and vinculin in *SACS* KO cells. c. Co-IP of NFASC and vinculin in WT and *SACS* KO cells. Vinculin was not detected in secondary antibody only control blots (not shown), suggesting a specific interaction. Quantification of n=3 co-IP experiments shows the interaction between VCL and NFASC is greatly reduced in *SACS* KO cells, despite NFASC being substantially overexpressed in *SACS* KO cells (Fig. 6d). d. STRING protein interaction map depicting proteins identified across proteomics datasets. Lines between proteins indicate high confidence interactions (interaction score>0.7). We removed proteins with redundant interactions for clarity (for example most integrins have largely overlapping interactomes). Proteins identified in the SACS interactome profiling are circled, with the thick circle marking interactors identified in all replicates, thin circle marking interactors identified in a subset of samples. Proteins are colored by log_2_ fold change in proteome (left half) and cell surface proteome (right half). Striped lines indicate no detection. Clusters identified by k-means clustering are marked by grey background.

To identify central proteins which may explain the cellular phenotypes in *SACS* KO cells, we performed STRING network analysis^49^. We considered all proteins altered in any dataset, and assessed only high confidence physical or regulatory interactions. K-means clustering of network interactions identified three clusters, each of which highlight complementary pathways by which sacsin contributes to cell structure and signaling (Fig. 7d). Central to cluster 1 is the interaction between sacsin and IF proteins, which interact with a variety of cell surface receptors. Combined with our biochemical experiments, this suggests that the loss of sacsin leads to improper localization of adhesion proteins to the plasma membrane, through decreased protein interactions between IFs, adaptors, and adhesion proteins. PTEN regulates several of these proteins, suggesting additional targets beyond FAK which may contribute to *SACS* KO phenotypes. The network also highlighted the microtubule associated kinase MAST1 (Fig. 1j, increased expression), which stabilizes PTEN^66^, and is protected from proteasomal degradation by the SACS interactor HSP90B1^67^.

Cluster 2 is composed of the interaction between sacsin, chaperone network proteins, and microtubules, which in concert regulate membrane protein processing, trafficking, and localization^68^. Multiple HSP chaperones were part of the sacsin interactome (Fig. 7d), including the marker of ER stress HSPA5/BIP, and several HSP90 proteins, which can stabilize FAK, modulate cell migration^69^, and regulate microtubules^70^. Recent evidence suggests HSP90 is essential for microtubule acetylation^71^, suggesting that the loss of sacsin may alter microtubule stability via HSP proteins (Fig. 2d,e). HSP proteins also regulate Rab proteins^72^ (cluster 3), which have diverse roles in vesicular trafficking, including PTEN and EGFR trafficking^73^. GO term analysis revealed that 65% of sacsin interacting proteins are involved in exosome related processes, with additional interactors being implicated in unfolded protein binding (HSPs) and focal adhesions (Extended Data Fig. 8c). In all, these results suggest that sacsin plays a critical role in functionally bridging protein quality control systems, microtubule dependent vesicular transport, and membrane localization of adhesion proteins.

## Discussion

This study identifies sacsin as a critical regulator of multiple aspects of cellular structure, including IF architecture, microtubules, protein trafficking, and focal adhesions. The complex and intertwined relationships between these processes complicates our understanding of their precise pathophysiological relevance, but our study raises some intriguing possibilities. Sacsin possesses a functional J domain, which interacts with HSP70 chaperone proteins^74,75^ (Fig. 7d). HSP proteins play a role in ubiquitin dependent turnover of IFs^76^, and neurofilament bundling in ARSACS neurons can be rescued by HSP expression^5^. Sacsin also possesses an ATPase domain with homology to HSP90 proteins. The sacsin interactor HSP90B1 stabilizes FAK^69^, suggesting that restoring FAK signaling may rescue IF structure through HSP activity (Fig. 4, 7d). It is also possible that sacsin transiently interacts with HSP90 regulated kinases, such as FAK^69^, and has a more direct role at focal adhesions. HSP70/90 complexes bind to microtubules in an acetylation dependent fashion^77^, and interact with hyperphosphorylated tau to increase tau’s interaction with microtubules^78^. Since HSPs are known to regulate all of the protein clusters with deficits in *SACS* KO cells (Fig. 7d), we hypothesize that the interaction between HSPs and sacsin may be an especially critical interaction that is lost in ARSACS. Furthermore, as illustrated by sacsin’s mediation of the interaction between IFs and focal adhesions, changes in additional as yet uncharacterized protein-protein interactions may explain specific ARSACS phenotypes, such as disrupted autophagy, nuclear morphology, and aberrant localization of mitochondria.

Why do mutations in sacsin, which is expressed throughout the brain, present as a cerebellar ataxia? Proteins whose abundance or localization are altered in *SACS* KO cells, and which also cause cerebellar ataxia, could suggest a causal molecular deficiency in ARSACS. The interactions between NFASC, NRCAM, and CNTN1 are critical for brain development, and mutation of each causes phenotypes reminiscent of ARSACS. *Cntn1* KO mice have deficits in axon guidance and develop cerebellar ataxia^79^. *Nrcam* KO mice have phenotypes only in lobules 4/5 of the cerebellar vermis^80^, which are also specifically affected in ARSACS^7,9,81^. Lastly, human mutations in *NFASC* which selectively remove the 155kD glial isoform cause congenital hypotonia, demyelinating neuropathy (as in ARSACS) and severe motor coordination defects^52^, while mutations of the neuron specific 186kD NFASC isoform cause cerebellar ataxia^53^. These convergent phenotypes lead us to hypothesize that improper localization of synaptic cell adhesion molecules is a causal molecular deficiency in ARSACS. Determining the mechanism by which sacsin controls the localization and function of these proteins throughout disease progression will be critical to our understanding of ARSACS pathophysiology, and developing rationally designed therapeutic strategies.

## ACKNOWLEDGEMENTS

We express our gratitude to Sonia Gobeil and the ARSACS patient community. This work was supported by grants to J.P.C and J.M.W. from the Fondation de l’Ataxie Charlevoix-Saguenay. We thank Thomas Sterns Karim Gilbert, Natalie Barker, and Dennis Goldfarb for technical assistance. We thank Dr. Stefan Strack for generously providing ARSACS mice. JPC was supported by the BBSRC (BB/R003335/1) and Ataxia UK. JMW was supported by grants from the National Institute for Child Health and Human Development (NICHD; T32HD040127) and a Pfizer-NCBiotech Postdoctoral Fellowship in Gene Therapy. LMG is supported by NIH R01 GM138520. The UNC Catalyst for Rare Diseases gratefully acknowledges the support of the Eshelman Institute for Innovation. This research is based in part upon work conducted using the UNC Proteomics Core Facility, which is supported in part by P30 CA016086 Cancer Center Core Support Grant to the UNC Lineberger Comprehensive Cancer Center. Confocal microscopy was performed using a microscope funded by Barts Charity (MGU0293). The UNC microscopy core is supported by the NICHD (P50HD103573).

## AUTHOR CONTRIBUTIONS

JMW and JPC conceived of the study. JMW, JPC, and LELR analyzed data and prepared the manuscript. SN, TMH, and JMW created *SACS* KO neuroblastoma lines. LMG aided in experimental design and provided reagents for proteomics experiments. AJH and KT provided reagents and experimental oversight. WYA, KMH, LX, and JMW performed histology. SH, TH, and WA managed the mouse colony. KMH, TMH, SH, JB, LEH, and JMW performed quantitative proteomics experiments. TVN and CLH performed RNAseq, KH and JMW analyzed the data. TVN and CLH performed the SACS interactome experiments. AH and KT performed co-IP MS experiments. LELR, WYA, KH, TVN, TMH, SH, LP, JMW, and GCA performed cell biology experiments.

---

**Extended Data Figure 1.**
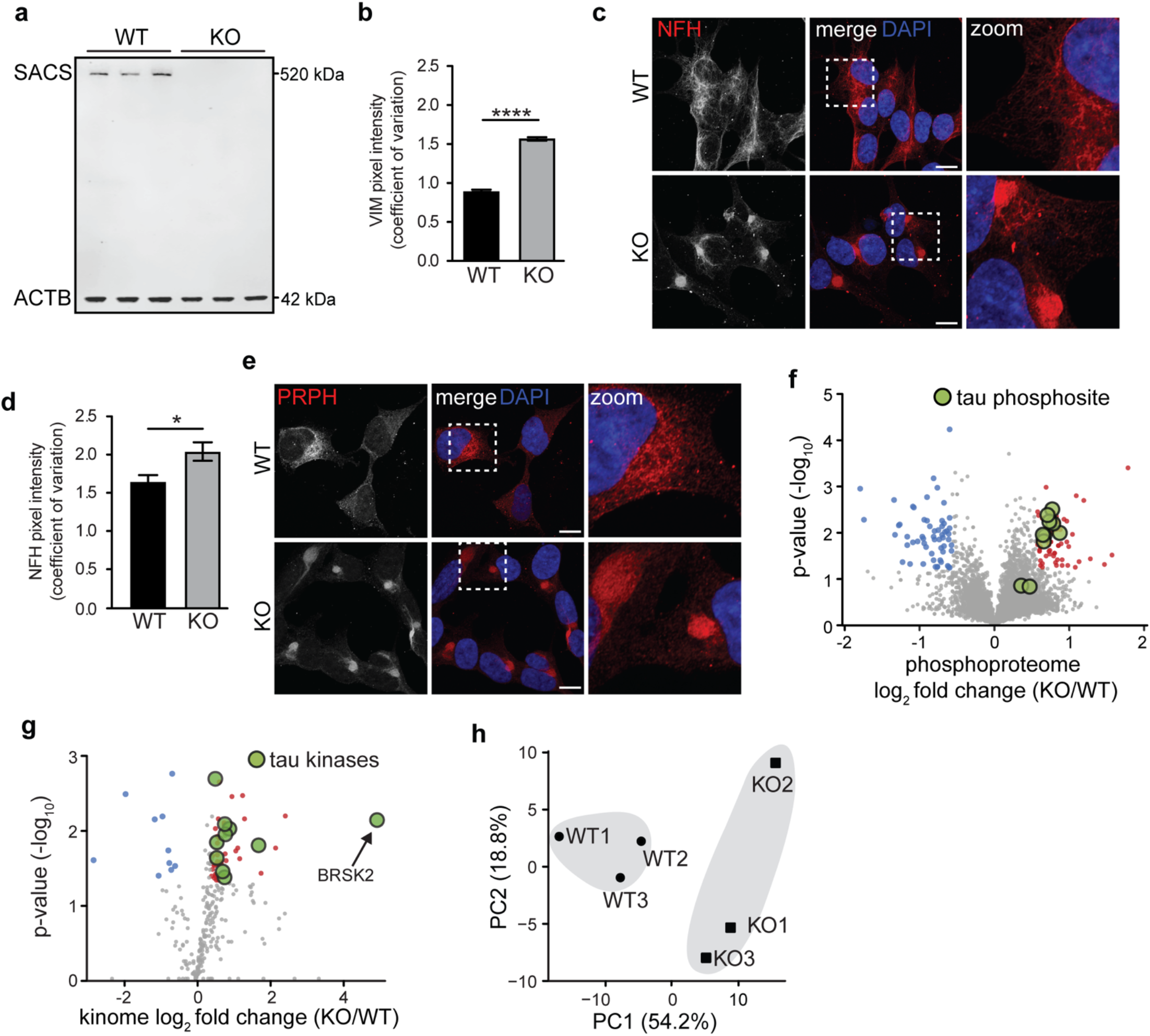
*SACS* KO SH-SY5Y cells recapitulate cellular phenotypes consistent with known deficits. a. Western blot for SACS and ACTB demonstrating the loss of SACS in SH-SY5Y KO cells. b. Coefficient of variation of vimentin pixel intensity values across the cell, with lower values indicating uniform distribution and higher values indicating polarized protein distributions. c. Representative confocal images of WT and *SACS* KO cells immunostained for the neurofilament heavy chain. d. Coefficient of variation of NFH pixel intensity. e. Representative confocal images of WT and *SACS* KO cells immunostained for peripherin, an intermediate filament protein found in neurons in the peripheral nervous system. f. Phosphoproteomic analysis of *SACS* KO SH-SY5Y cells. Green circles mark specific phosphorylated residues on tau. g. Kinome profiling of *SACS* KO SH-SY5Y cells. Green circles mark kinases which are known to directly phosphorylate tau. h. Principle component analysis of all kinases identified in kinome profiling data (Supp. Table 1). Unsupervised hierarchical clustering separated WT and KO cells (grey shading), suggesting widespread changes in the kinome of *SACS* KO cells.

**Extended Data Figure 2.**
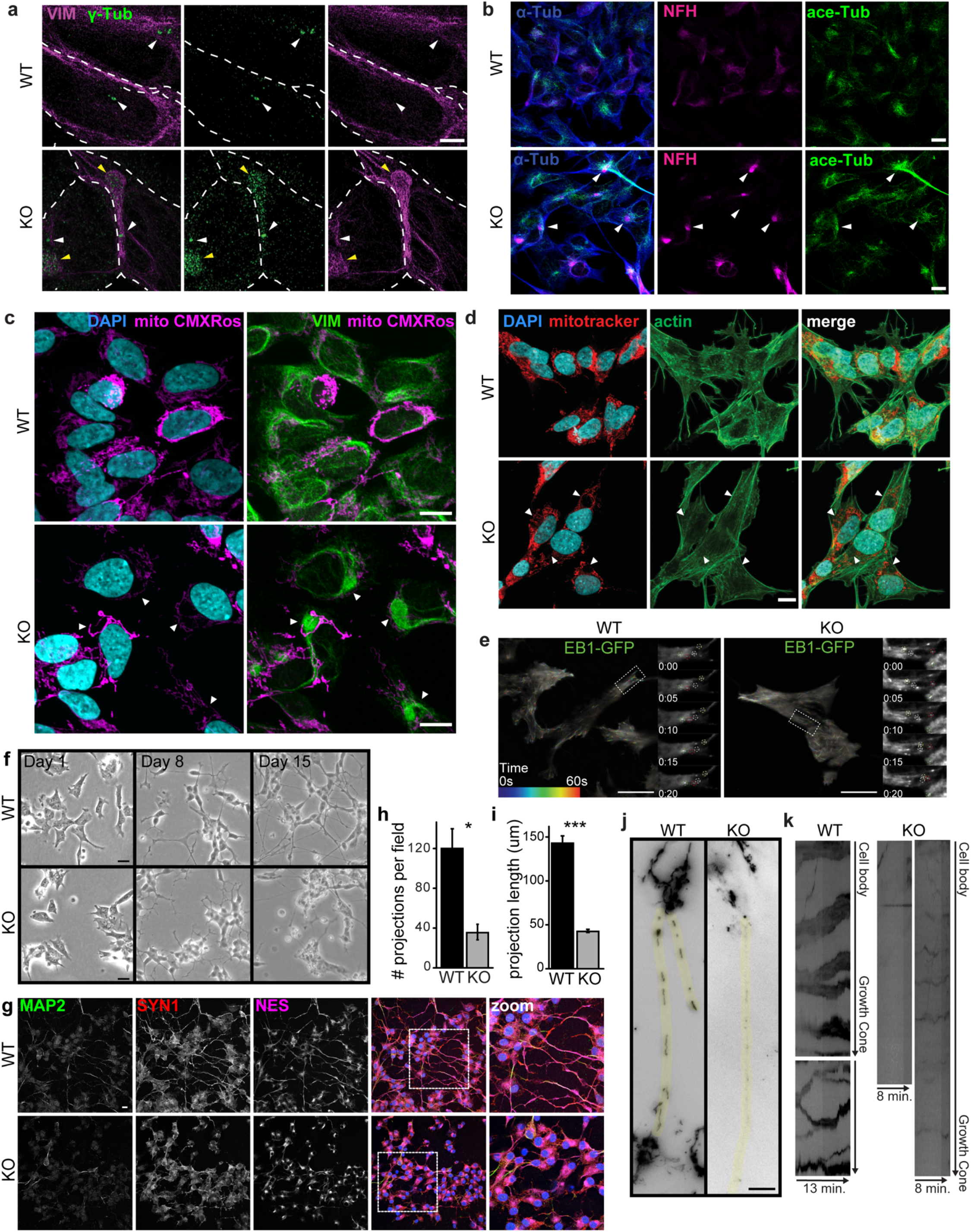
Microtubule and mitochondria deficits in *SACS* KO cells. a. Super resolution structural illumination microscopy images showing accumulation of gamma-tubulin within perinuclear vimentin bundles of *SACS* KO cells. White arrows point to centrioles, yellow arrowheads highlight the presence of gamma-tubulin within vimentin bundles in KO cells. Dashed white lines denote boundaries between adjacent cells. Scale bar = 1um. b. Representative confocal images of immunostaining for alpha tubulin, neurofilament heavy, and acetylated tubulin in WT and *SACS* KO cells. Arrowheads mark coincidence of acetylated tubulin and neurofilament bundles, suggesting that acetylated tubulin structures are found in proximity to neurofilament bundles, but also localize throughout the cell. c. Representative confocal images of WT and *SACS* KO cells stained for the mitochondria membrane potential dependent dye CMXRos, vimentin, and nuclei (DAPI). Arrowheads highlight the exclusion of mitochondria from vimentin bundles. d. Representative confocal images of WT and *SACS* KO cells immunostained for mitotracker, actin, and nuclei (DAPI). Arrowheads highlight the exclusion of mitochondria from vimentin bundles. e. Representative TIRF microscopy images from WT and *SACS* KO cells expressing EB1-GFP. Microtubule growth tracks are color coded marking their position over time. Insets show the enlargement of outlined regions and movement of individual comet movement over time (circles), numbers refer to seconds. f. Representative phase contrast brightfield images of WT and *SACS* KO cells across 15 days of neuronal differentiation. g,h. Quantitation of the number of projections per field (g) and length of projection (h) of WT/KO cells demonstrating significantly reduced number and length of projections in *SACS* KO cells. i. Confocal images of WT/KO cells after 15 days in differentiation conditions, stained for neuronal markers microtubule associated protein 2 (MAP2) and synapsin 1 (SYN1), and the intermediate filament protein nestin (NES), a marker of immature neurons. Scale bar = 10 μm. j. Mitochondria labeled with mitoTracker GreenFM in neurites (highlighted in yellow) of 15 day differentiated WT/KO cells demonstrating the lack of elongated mitochondria in *SACS* KO neurites. Images were snapshots from live-cell time-lapse imaging. k. Kymograph illustrating mitochondrial transport along neurites of differentiated WT/KO cells. Note that mitochondrial undergo both retrograde and anterograde movement in control but are relatively static in *SACS* KO cells. Scale bars = 10 μm.

**Extended Data Figure 3.**
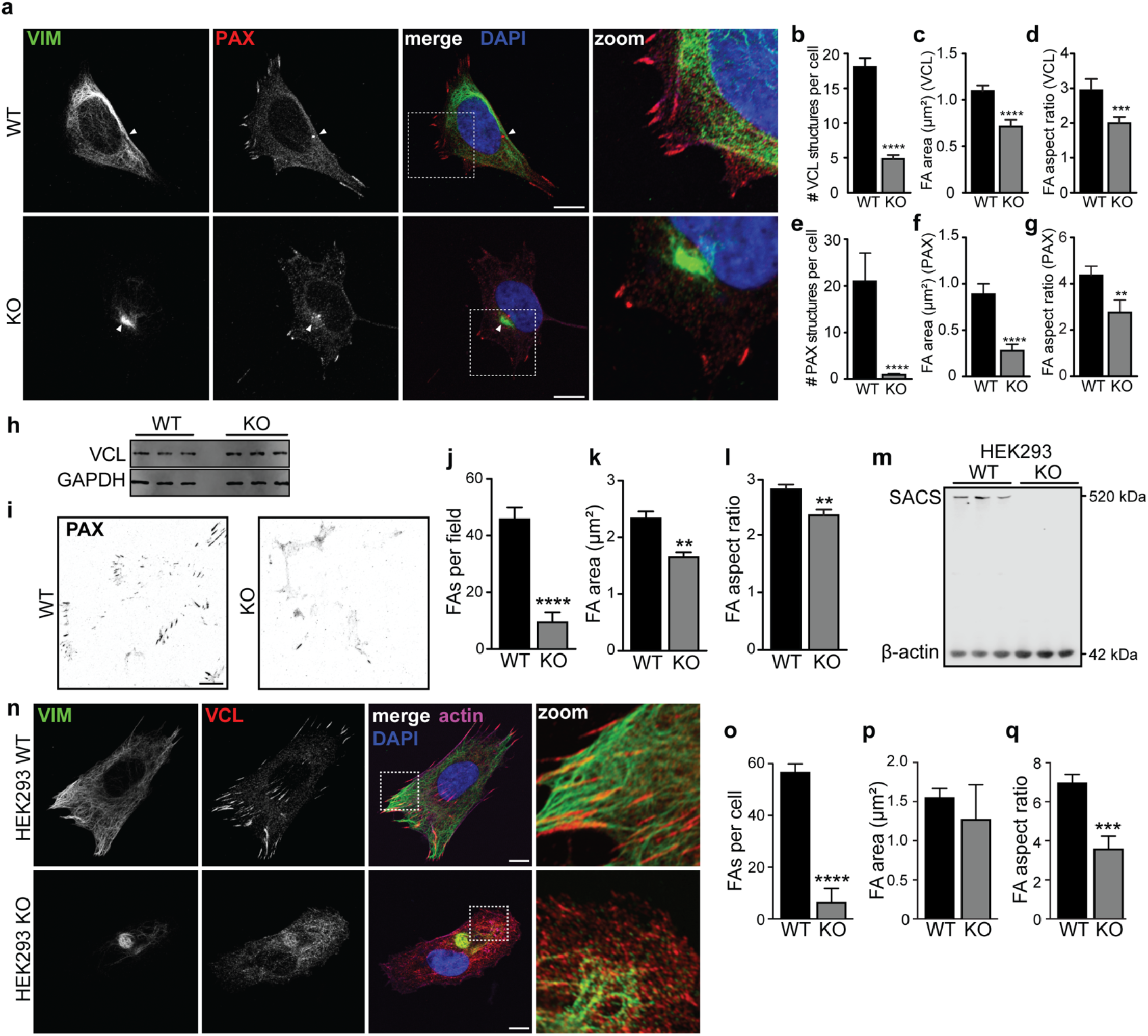
Focal adhesions are disrupted in *SACS* KO cells. a. Representative confocal image of WT/KO cells labelled with vimentin and paxillin. Arrowhead marks the PAX positive MTOC, which is sequestered in the vimentin bundle in *SACS* KO cells. b-g. Quantification of images from Fig. 3c (b-d), and Extended Data Figure 3a (e-g). Aspect ratio = width:height ratio. n=3 independent replicates. h. Western blot for vinculin, showing that levels of the focal adhesion protein are unaltered in KO cells. i. Representative image of cover slips treated with hypotonic shock to remove cell bodies, leaving focal adhesions retained through ECM interaction. j-l. Quantification of the incidence, area, and aspect ratio of paxillin positive focal adhesions in WT/KO cells. m. Western blot for SACS and ACTB demonstrating the loss of SACS in HEK293 KO cells. n. Confocal images of HEK293 cells immunolabeled for vimentin, vinculin, and actin. o-q. Quantification of images from Supp. Fig. 3n, suggesting focal adhesion deficits are consistent with SH-SY5Y cells. r. Representative confocal image of WT/KO SH-SY5Y cells immunolabelled for ITGA6 and ITGB1.

**Extended Data Figure 4.**
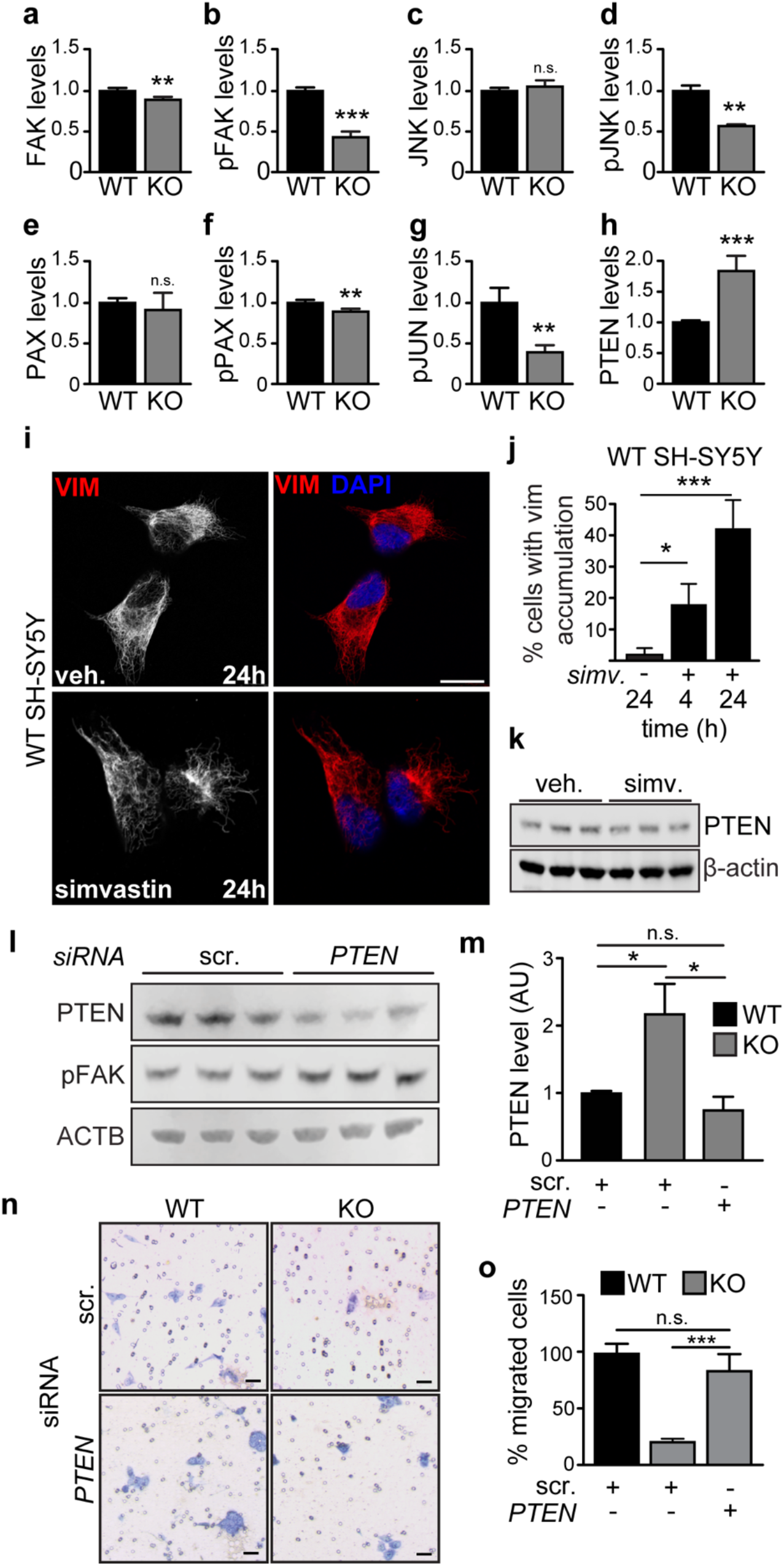
Modulating *PTEN* rescues cellular phenotypes in *SACS* KO cells. a-h. Quantification of immunoblots from Fig. 4c. Intensity normalized to ACTB. n=3 biological replicates. i. Representative confocal images of the induction of vimentin bundling by simvastin. Scale bars = 10 μm. j. Quantification of vimentin bundling phenotype induced by simvastin over time. k. Western blot of PTEN levels in 24 hour simvastin treated WT cells, suggesting that vimentin bundling does not affect PTEN levels. l. Representative Western blot of *SACS* KO cells treated with siRNAs targeting *PTEN* or scrambled. m. Quantification of PTEN levels in WT/KO cells treated with scrambled or *PTEN* targeting siRNAs, suggesting *PTEN* is returned to WT levels in *SACS* KO cells. n=3. n. Representative images of WT/KO cells plated in Transwell chambers with 8 μm pores, allowed 24 hours for cells to migrate through pores, and fixed and stained with Giemsa blue. Scale bar = 20 μm. o. Quantification of transwell assays in WT/KO cells treated with the indicated siRNAs. n=5 per cell line.

**Extended Data Figure 5.**
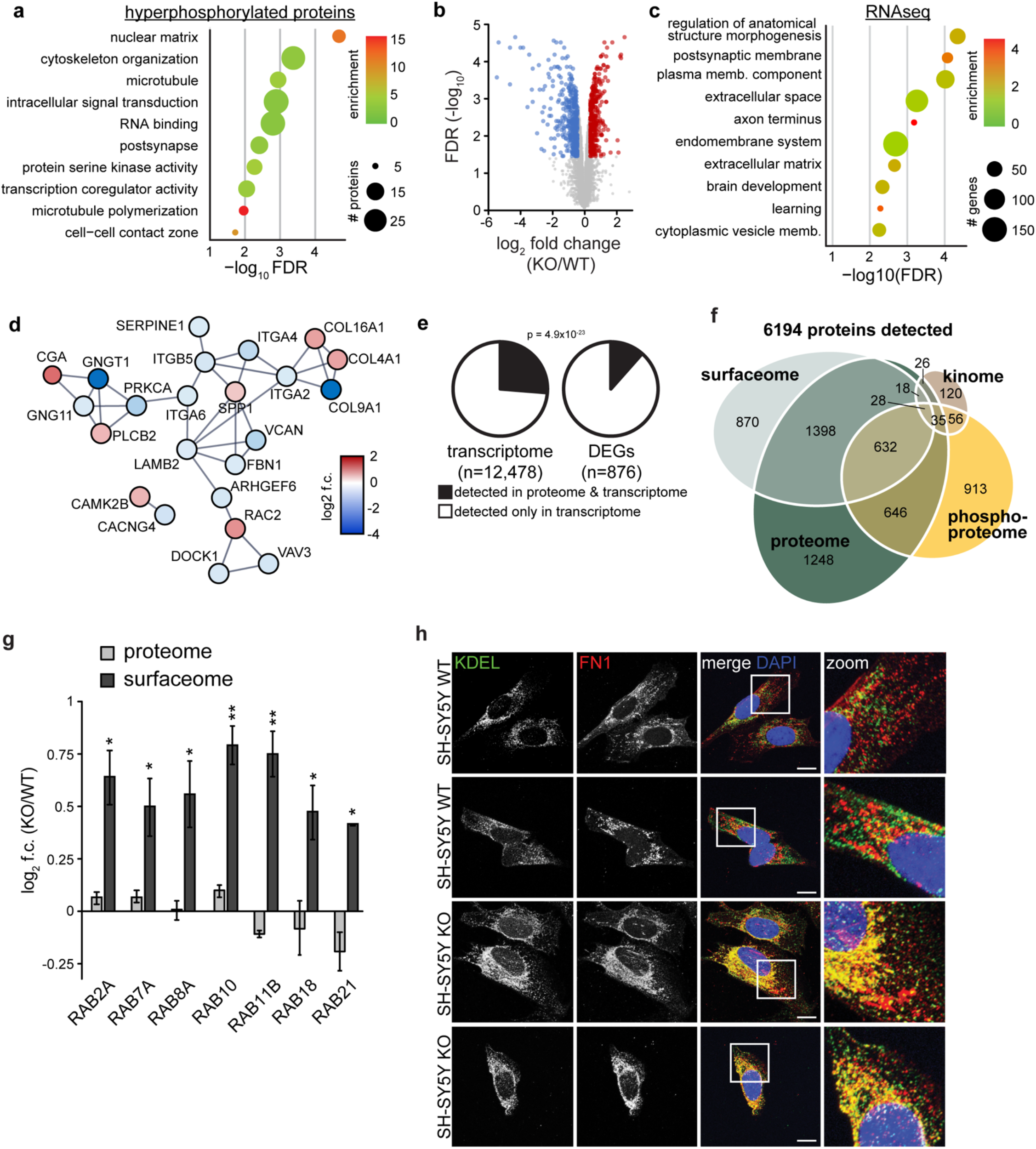
Altered transcription of synaptic adhesion and vesicular proteins. a. GO term analysis of hyperphosphorylated proteins in proteomic analysis (p<0.05, log_2_ fold change -/+ 0.4) (Extended Data Fig. 1f), identifying multiple terms related to nuclear/transcriptional processes. b. RNAseq of 15 day neuronally differentiated SH-SY5Y cells. c. GO term analysis of differentially expressed genes suggests that synaptic and vesicular transport genes are altered in neurons (p<0.05, log_2_ fold change -/+ 0.5). d. Interaction network of cell adhesion proteins that are differentially expressed. e. Overlapping gene/protein identification from RNAseq and proteomics, showing that DEGs were not detected as readily as genes in the rest of the transcriptome. Hypergeometric test was used to calculate enrichment p-value. f. Euler diagram of protein identification across all mass-spec datasets. g. Log_2_ fold change of Rab proteins in proteome and surfaceome datasets. Asterisks refer to statistical significance in each dataset. No Rabs were significant in the proteome. h. Representative confocal images of cells immunolabelled for fibronectin and KDEL in in WT/KO SH-SY5Y cells.

**Extended Data Figure 6.**
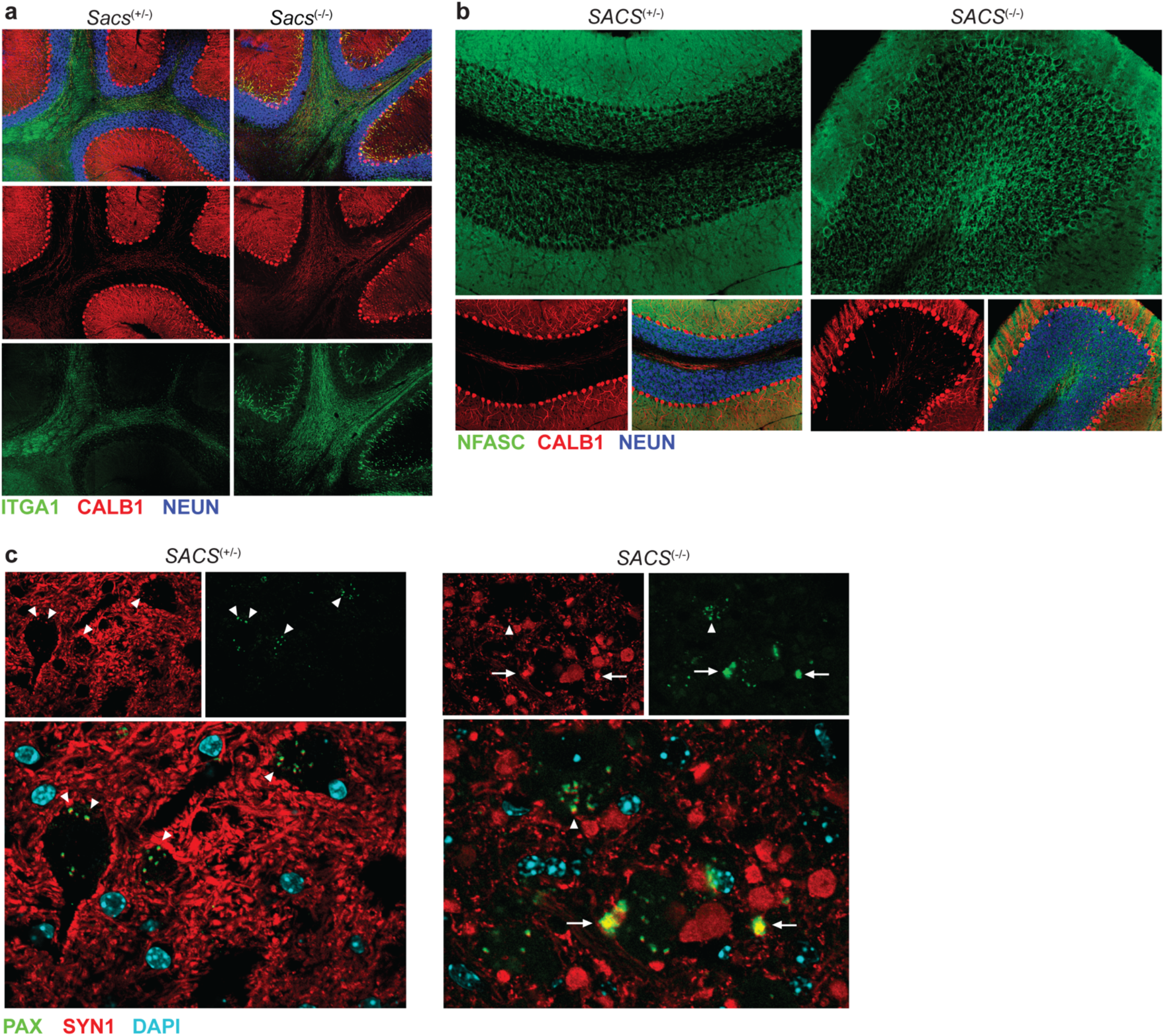
Cerebellar imaging in *SACS* KO mice. a. Widefield view of ITGA1 staining in *Sacs*^(+/-)^ and *Sacs^(-/-)^* mice. ITGA1 staining is normally observed along Purkinje neuron axon tracts, but is mislocalized to the proximal dendritic arbor in *Sacs^(-/-)^* mice. b. Widefield view of NFASC expression in dorsal cerebellar lobules in P120 mice, demonstrating increased NFASC intensity surrounding Purkinje neuron soma in *Sacs*^(-/-)^ mice. c. Representative confocal image of neurons (black voids) in the DCN. Arrowheads mark colocalization between paxillin and synapsin, arrows mark large extracellular PAX+/SYN+ aggregates.

**Extended Data Figure 7.**
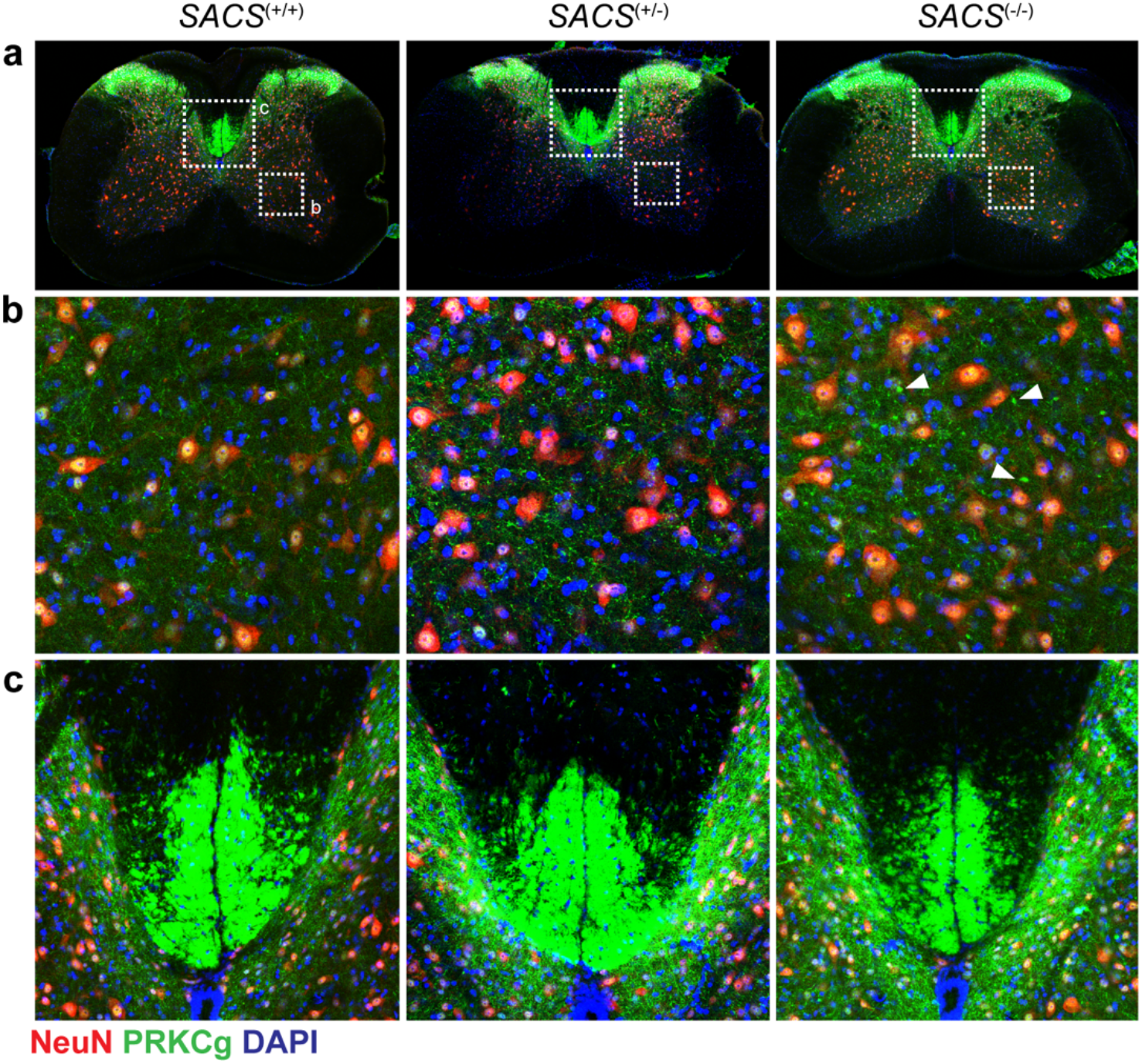
Axonal and synaptic alterations in spinal cord from P120 *SACS* KO mice. a. Representative confocal images of the lumbar spinal cord labeled for PRKCg (a neuronal marker specific to cortical layer 5 neurons^82^) NEUN, and DAPI in WT, *SACS*^(+/-)^, and *SACS*^(-/-)^ mice. b. High magnification of area of ventral spinal cord indicated in panel a. White arrowheads mark abnormal axonal swellings, similar in appearance to those surrounding DCN neurons. c. High magnification of lumbar the corticospinal tract indicated in panel a, showing an overall decrease in axonal tract density in *SACS*^(-/-)^ mice.

**Extended Data Figure 8.**
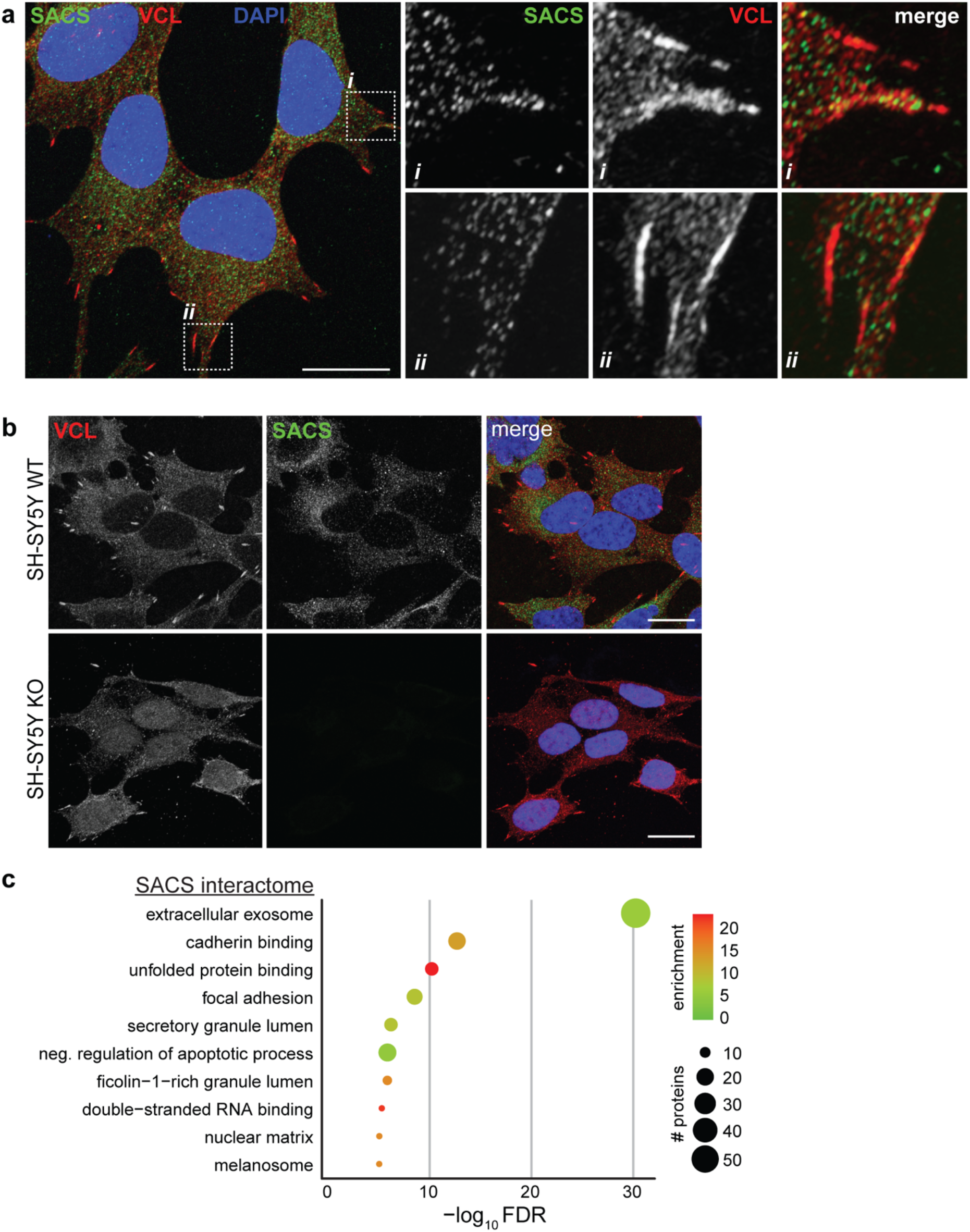
SACS interactors. a. Representative confocal image for sacsin and vinculin in WT SH-SY5Y cells demonstrating SACS colocalizes with focal adhesions. b. Representative confocal image for *SACS* KO cells processed in parallel to (a), demonstrating the specificity of SACS staining. c. GO term analysis of all proteins identified in the SACS co-IP interactome (Supplementary Table 4).

