## Supplementary material for "The ataxia protein sacsin is required for integrin trafficking and synaptic organization": Materials and Methods

#### Antibodies

All antibodies used in this study are listed in Supplementary Table 5.

#### Genome editing to generate sacsin KO SH-SY5Y cells

Human SH-SY5Y neuroblastoma cells were obtained from the American Type Culture Collection and were grown in 1:1 Dulbecco's Minimum Eagle Medium (DMEM)/Ham's F12 medium, plus 10% heat-inactivated foetal bovine serum,100 U $ml^{-1}$ penicillin and 100 mg $ml^{-1}$ streptomycin. An SH-SY5Y cell line with the sacsin truncation mutation M783* was generated using CRISPR/Cas9. We cloned the *SACS* targeting guide RNA (gRNA) TTTCATGGCTTAAGATGGTTTGG (PAM sequence underlined) into the p1261_GERETY_U6_BasI_gRNA vector for expression of the gRNA under control of the U6 promoter. The gRNA expression vector was co-transfected with a Cas9 expression vector (hCas9, Addgene # 41815) and a targeting vector with homology arms to introduce the M783* mutation along with a puromycin selection cassette (pMCS-*SACS*$^{trunc}$-PB:*PGKpuroDtk*) using Lipofectamine 3000. Puromycin-resistant clones were selected and screened by PCR and sequencing.

#### Proteomic, phosphoproteomic sample preparation

SH-SY5Y were grown to ~80% confluency in 13 T75 flasks for harvesting. SH-SY5Y flasks were placed on ice, washed twice with ice cold PBS and harvested by scraping cells in lysis buffer (50 mM HEPES, pH 7.5, 150 mM NaCl, 0.5 % Triton X-100, 1mM EDTA, 1mM EGTA, 10mM NaF, 2.5mM Na3VO4, complete protease inhibitor cocktail (Roche), and phosphatase inhibitor cocktail 2 and 3 (Sigma). Lysates were sonicated by pulsing once at 30% for 10 seconds and then twice at 40% for 10 seconds with 10 second rest on ice between each pulse (Branson 150 Sonifier). Lysate was transferred to microcentrifuge tube and spun at 14,000 x g for 10 minutes at 4°C. Lysate was filtered through a 0.2 µm syringe filter and stored at -80°C until all replicates were collected. Protein concentration was quantified using a Bradford assay. Cell lysates (1mg, n=3) were acetone precipitated overnight and stored at -20C. Protein pellets were resuspended in 8M urea, then reduced with 5M DTT at 56C for 30 min and alkylated with 15mM iodoacetamide in the dark at RT for 45 min. Samples were diluted to 1M urea, then digested overnight with trypsin (Promega) at 1:100 trypsin:protein ratio. Samples were acidified then desalted using C18 desalting spin columns (Pierce). A peptide BCA colorimetric assay (Pierce) was performed and 500µg of each sample was individually labeled with TMT6 reagent (Thermo). After labeling efficiency was confirmed, the TMT6 labeled samples were mixed and desalted using C18 desalting spin column (Pierce). A 100ug aliquot was set aside for global proteome analysis, and was fractionated into 4 fractions using a High pH reversed phase fractionation spin column (Pierce). The rest of the sample (~3mg) was enriched for phosphopeptides using Ti-MAC magnetic beads (ReSyn Biosciences). The Ti-MAC eluate was fractionated into 3 fractions using High pH reversed phase fractionation spin column.

Global proteome and phosphoproteome fractions were dried down via vacuum centrifugation and stored at -80C until LC/MS/MS analysis.

**MIB/MS Kinome analysis**

To enrich for kinases, multiplexed inhibitor bead (MIB) kinase enrichment was performed as previously described with a slightly modified bead composition[83]. Specifically, each sample was applied to an individual 350 µL Poly-Prep® chromatography column (Bio-Rad) containing the following immobilized kinase inhibitors (CTx-0294885, PP58, Purvalanol B, UNC2147A, VI-16832, UNC8088A). Kinase elution, removal of detergents and isolation of peptides was completed as described before[83]. Kinase peptides were desalted and submitted to the UNC Michael Hooker Proteomics Core for LC/MS/MS analysis.

**Cell surface labeling**

For cell surface labelling, $2x10^6$ SH-SY5Y WT and SACS KO cells were each plated in nine 10 cm dishes, and cultured until 95% confluent (~3 days) (n=3 per cell line). To identify proteins which purify non-specifically an additional replicate of WT/KO lines were processed as below, but without the addition of Biocytin hydrazide. Cells were lifted using CellStripper Dissociation Reagent (Corning #25056CI) for 20 minutes at 37°C and resuspended in 1X PBS (pH 6.5) + 1.6 mM NaIO4 and rotated at 4°C for 20 minutes in the dark. Cells were washed three times then resuspended in 1X PBS (pH 6.5) + 10mM Aniline + 1mM Biocytin hydrazide and incubated at room temperature for 60 minutes, then at 4°C for 20 minutes while rotating. After three PBS washes, cell pellets were resuspended in RIPA, rotated at 4°C for 30 minutes, and sonicated with 1 second pulses at 20% power for 1 minute. To enrich for the labeled surface proteins, cells were centrifuged 15,000 rpm for 10 minutes at 4°C and supernatant was incubated in washed Neutravidin High-Capacity Resin (ThermoFisher #29204) for one hour at 4°C. Resin was added to gravity column and washed with RIPA, 1X PBS (pH 7.4) + 1M NaCl, Ammonium Bicarbonate (ABC) + 2M Urea then resuspended in ABC + 2M Urea + 5 mM tris(2-carboxyethyl)phosphine (TCEP) and incubated at room temperature in the dark at 55°C shaking at 300 rpm for 30 minutes. Iodoacetamide (IAM) was then added to a final concentration of 11 mM and shaken at room temp for 30 minutes in the dark. Resin was centrifuged at 500g for 5 minutes and resuspended in 1 ml ABC + 2M urea containing 20 ug trypsin (Fisher #P8101) to fragment peptides at RT overnight. To desalt, samples were by acidified to pH<2 with 10% trifluoroacetic acid (TFA) in C-18 spin column (ThermoFisher #89873), washed, and resuspended in 40% acetonitrile + 0.1% formic acid then dried with vacuum centrifugation and stored at -80˚ C.

**LC-MS/MS Analysis**

Kinome, proteome and phosphoproteome samples were analyzed by LC/MS/MS using an Easy nLC 1200 coupled to a QExactive HF mass spectrometer (Thermo Scientific). Samples were injected onto an Easy Spray PepMap C18 column (75 µm id × 25 cm, 2 µm particle size) (Thermo Scientific) using a 120 min method. The gradient for separation consisted of 5–50% mobile phase B at a 250 nl/min flow rate, where mobile phase A was 0.1% formic acid in water and mobile phase B consisted of 0.1% formic

acid in 80% ACN. The QExactive HF was operated in data-dependent mode where the 15 most intense precursors were selected for subsequent HCD fragmentation.
For kinome samples, QExactive HF was operated as previously described[83]. QExactive HF resolution for the precursor scan (m/z 350–1600) was set to 120,000 with a target value of $3 \times 10^6$ ions and a maximum injection time of 100 ms. MS/MS scans resolution was set to 60,000 with a target value of $1 \times 10^5$ ions and a maximum injection time of 100 ms. The normalized collision energy was set to 27% for HCD with an isolation window of 1.6 m/z. Dynamic exclusion was set to 30 s, peptide match was set to preferred, and precursors with unknown charge or a charge state of 1 and ≥ 8 were excluded.
For TMT proteome and phosphoproteome samples (each biological replicate analyzed in duplicate), QExactive HF resolution for the precursor scan (m/z 350–1600) was set to 60,000 with a target value of $3 \times 10^6$ ions and a maximum injection time of 100 ms. MS/MS scans resolution was set to 60,000 with a target value of $1 \times 10^5$ ions and a maximum injection time of 100 ms. Fixed first mass was set to 110 m/z and the normalized collision energy was set to 32% for HCD with an isolation window of 1.2 m/z. Dynamic exclusion was set to 30 s, peptide match was set to preferred, and precursors with unknown charge or a charge state of 1 and ≥ 8 were excluded.
Cell surface samples were analyzed by LC-MS/MS using a Thermo Easy nLC 1200 coupled to a Thermo Fusion Lumos mass spectrometer. Samples were injected onto a Thermo PepMap C18 trap column, washed, then loaded onto an Easy Spray PepMap C18 analytical column (75 μm id × 25 cm, 2 μm particle size) (ThermoFisher). The samples were separated over a 120 min method, where the gradient for separation consisted of 5–45% mobile phase B at a 250 nl/min flow rate; mobile phase A was 0.1% formic acid in water and mobile phase B consisted of 0.1% formic acid in 80% acetonitrile. MS1 orbitrap scans were collected at a resolution of 120,000 and 1e6 AGC target. The MS2 scans were acquired in the Orbitrap at 15,000 resolution, with a 1.25e5 AGC, and 50ms maximum injection using HCD fragmentation with a normalized energy of 30%. Dynamic exclusion was set to 30 seconds and precursors with unknown charge or a charge state of 1 and ≥ 8 were excluded.

**Kinome, proteome, phosphoproteome, and cell surface data analysis**
Kinome, proteome and phosphoproteome raw data files were MaxQuant version 1.6.1.0 and searched against the reviewed human database (downloaded Feb 2017, containing 20,162 entries), using Andromeda within MaxQuant. Enzyme specificity was set to trypsin, up to two missed cleavage sites were allowed, carbamidomethylation of C was set as a fixed modification and oxidation of M and acetyl of N-term were set as variable modifications. For phosphoproteome samples, phosphorylation of S,T,Y was set as a variable modification. For proteome and phosphoproteome samples, the quantitation type was set to reporter ion MS2 and for kinome label-free quantitation, match between runs was enabled. A 1% FDR was used to filter all data. For kinome data, a minimum of two peptides was required for label-free quantitation using the LFQ intensities.
Cell surface proteome raw data files were processed using MaxQuant version 1.6.15.0 and searched against the reviewed human database (downloaded Feb 2020, containing 20,350 entries), using Andromeda within MaxQuant. Enzyme specificity was set to trypsin, up to two missed cleavage sites were allowed, carbamidomethylation of C was

set as a fixed modification and oxidation of M and acetyl of N-term were set as variable modifications. A 1% FDR was used to filter all data and match between runs was enabled. A minimum of two peptides was required for label-free quantitation using the LFQ intensities.

For all proteomic datasets, proteins with a missing value in one replicate were imputed using the KNN imputation method, proteins with two or more missing values were removed from analysis. Linear Models for Microarray Data (LIMMA) was used to calculate log2 fold change of LFQ intensity and perform statistical analysis[84]. For proteins identified in the surfaceome, we annotated them as 'Membrane" or 'Exosome' based on DAVID bioinformatics database. Proteins which were identified in unlabeled controls (no biotin) were removed from further analysis. Proteins with $p<0.05$ and log2 fold change of KO/WT -/+>0.4 were included in downstream analyses. The Kinome tree was generated on CORAL[85].

### GO Term Analysis

For GO term graphs, the list of significant genes for each proteomic experiment were input into geneontology.com with human genome as the background, Fisher's exact test with FDR correction[86]. Graphs include terms in all categories (biological processes, molecular function, cellular component). Due to the hierarchical nature of GO terms in Panther (i.e. groups of terms have a nested nature to assign relationships between them) we only considered the most proximal term in each hierarchy to ensure terms were specific and directly comparable. These proximal terms are listed as "PARENT" in Supplementary Table 2. All terms underneath each parent are also listed for reference. Terms were ranked by FDR value and the top ten non-redundant top terms were included in each figure. All terms, including graphs were generated using R scripts from bioprotocol.org (DOI: 10.21769/BioProtoc.3429).

### Western Blots

Cells were lysed with RIPA buffer and 20 ug protein was loaded per lane and on a Novex 4-12% (4-20% for BRSK2) Tris-Glycine gel (ThermoFisher). Protein was transferred to PVDF and blocked with 1X Blocker BSA (ThermoFisher). Blots were washed and incubated with HRP secondary antibodies (ThermoFisher). Protein was quantified using ImageJ software. Each lane was normalized to the relative density of GADPH/ACTB.

Cell surface protein isolation (Fig. 5a) was validated through western blot by collecting 250 ul resin (bound to cell surface proteins) from the wash column and centrifuging at 2000xg for 5 minutes to pellet resin and remove residual 50mM ABC + 2M Urea. Resin was then resuspended in 250 ul 2X Laemmli buffer with 5% 2-mercaptoethanol. The control sample contained 10 ul of total cell lysates (collected prior to resin addition) from SH-SY5Y WT and SACS KO cells. The samples were run on a 4-15% TGX Bio-Rad pre-stained gel (Bio-Rad #4568094) and transferred to a PVDF membrane, which was blocked in 5% milk in 1X TBST. Primary antibodies were diluted 1:1000 in 5% BSA in 1X TBST. Goat anti-rabbit HRP secondary antibody was diluted 1:5000 in 5% milk. Remaining immunoblotting was as described previously[4].

### Fractionated Western Blots

SH-SY5Y WT and SACS KO cells were fractionated with the Minute Plasma Membrane Protein Isolation and Cell Fractionation Kit (Invent). The cytoplasmic fraction was lysed in Buffer A and the plasma membrane fraction was lysed in RIPA buffer, both fractions contained protease inhibitor (Pierce). Proteins were diluted 1:1 with 2x laemmli buffer with betamercaptoethanol and run on TGX prestained gels (Biorad). Total protein images were obtained before transferring onto a PVDF membrane. Individual protein levels were normalized to total protein image using Image J software for quantification.

**Endogenous Co-immunoprecipitation**

SH-SY5Y cells were subjected to chemical cross-linking. The cleavable, homo-bifunctional cross-linker dithiobis[succinimidylpropionate] (DSP; Pierce, Rockford, IL, USA) was diluted to a final concentration of 1mM in PBS and added to the cultured cells. After incubation for 1h at room temperature, cross-linking was stopped by addition of Tris (pH 7.5) to a final concentration of 20mM. Cells were then washed twice in ice-cold PBS, before the cells were harvested in RIPA buffer (NFASC IP) or 50mM Tris-HCl pH7.4 + 150mM NaCl + 1mM EDTA + 0.5% Triton X-100, supplemented with protease inhibitors (Pierce/Roche) and incubated on ice for 5 minutes (SACS/VIM IP).
Sacsin and vimentin IP: A small aliquot of supernatant was removed for analysis by immunoblotting (input fraction) the remaining supernatant was incubated with rabbit monoclonal anti-vimentin antibody or rabbit monoclonal anti-sacsin overnight at 4°C on a rotor. After 16 hours 50µl of magnetic beads (Sigma, Poole, UK) were washed twice in PBS-Tween 0.1% buffer, before being recovered in a magnetic separator. The beads were then resuspended within the cell lysate already incubated with the antibody for 2h at room temperature and washed.
NFASC IP: Protein concentration was assessed with a Bradford assay and a small aliquot of supernatant was removed for analysis by immunoblotting (input fraction). 300 ug of the sample were used for the immunoprecipitation. First, 20 ul of protein A magnetic beads (73778, CST) washed with 1x Cell Lysis buffer (9803, CST) were added to the samples for 20 minutes for preclearing. Next, beads were removed and 5 ul of the precipitation antibody, anti-neurofascin (PA5-78668, Invitrogen) or normal rabbit IgG isotype control (2729, CST), were added to the samples and incubated overnight at 4°C on a rotor. After 16 hours, fresh prewashed magnetic beads were added to the samples and rotated at room temperature for 20 minutes before being recovered in a magnetic separator and washed five times with 1x Cell Lysis Buffer.

**Cell culture immunostaining image acquisitions**

All cell culture samples, with the exception of samples prepared for microtubule staining, were fixed with 4% PFA and permeabilized with 0.1 to 0.3% Triton X-100. For microtubule staining, samples were fixed with 100% methanol for 20 minutes at -20 °C. Non-specific binding was reduced by blocking in 5% Normal donkey serum or 1% bovine serum albumin (BSA)+10% normal goat serum. Cells were incubated with primary antibodies for a minimum of 1 hour at RT, followed by incubation with labelled secondary antibodies, Phalloidin, and Hoechst for 1 hour at RT. Samples were mounted either in Fluoro-Gel mounting medium (Electron Microscopy Sciences, Cat. # 17985-30) or ProLong Diamond Antifade Mountant (Invitrogen, Cat. # P36961). Widefield images were captured with GE IN Cell 2200 high content imaging system equipped with a Plan

Fluor 20x/0.75 NA air objective. Confocal images were acquired using an inverted Olympus FV3000RS. Plan Apo 60x/1.4 NA oil (Olympus) or Plan Apo 30x/1.05 NA silicon oil (Olympus) objectives were used. SIM images were acquired and reconstructed in 3D-SIM mode using a Nikon N-SIM system equipped with a Plan Apo TIRF 100x/1.49 NA oil objective.
For focal adhesion immunolabelling, coverslips were coated with 10μg/ml fibronectin solutions overnight. Confocal microscopy was performed using a LSM880 (Zeiss) with a 63x objective and an AiryScan module. Quantification of incidence of cells with perinuclear vimentin accumulation and incidence of focal adhesion was performed blind to experimental status. Imaging processing was carried out with Zen Blue software (Zeiss). Focal adhesion isolation from cells was performed by hypotonic shock to remove cells while leaving focal adhesions, as described previously[87]. Isolated focal adhesions were then immunolabelled to detect vinculin and analysed using confocal microscope with quantification as described previously. All image processing steps were carried out using ImageJ software.

**EB1-GFP imaging, tracking and quantification**
For Total Internal Reflection Fluorescence (TIRF) imaging, SH-SY5Y cells were cultured on 35 mm glass bottom dishes (Mattek) and transfected with plasmid expressing EB1-GFP using FuGene transfection reagent (Promega). pGFP-EB1 was a gift from Lynne Cassimeris (Addgene plasmid #17234). TIRF live cell imaging was carried out 24 hours after transfection on an inverted Nikon Eclipse Ti2 equipped with a Plan Apo TIRF 100x/1.49 NA oil (Nikon) objective. TIRF images were captured at single z-plane, every second for a period of 1 minute. Tracking of EB1-GFP comet was performed using the FIJI plugin TrackMate with the following analysis settings: Laplacian of Gaussian detector with an estimated spot diameter of 0.16 μm, subpixel localization enabled, simple LAP tracker, minimum number of spots on track >9, and maximum number of spots on track <35. Each trajectory was visually inspected to confirm tracking accuracy. Mean track velocities were plotted and statistical analyses were performed using GraphPad Prism 9.0.

**Nocodazole experiment**
Cells were treated with 10 μm nocodazole (Sigma, Cat. #SML1665) or DMSO control for 3 hours at 37 °C. Washout of nocodazole was performed by one wash with ice old PBS and one wash with pre-warmed media. Cells were fixed with 100% methanol at indicated timepoints. Measurement of microtubule staining in untreated cells and repolymerization after nocodazole washout were performed on images constructed with maximum intensity projections. Areas occupied by microtubules and cells were thresholded and quantified using a custom CellProfiler pipeline. Microtubule density was quantified by thresholding and normalizing area occupied by microtubules to cell area.

**SH-SY5Y Differentiation**
SH-SY5Y cells were differentiated using a modified version of a previously described method[34]. Briefly, cells were grown to near confluence under normal maintenance conditions prior to the start of differentiation. $1X10^5$ cells/well were seeded in a 6 well plate. After 24 hours media was changed to DMEM containing 2% Fetal Bovine Serum

(FBS: Gibco, #A3840001) and10 µM of all-trans retinoic acid (ATRA: Sigma, #R2625). At day 4 cells were passaged and resuspended in neurobasal media (Gibco, #21103049) supplemented with B27 (Gibco, #17504044) 50 ng/ml Brain-Derived Neurotrophic Factor (BDNF: Sigma, #B3795), and 10 µM RA. At day 12 media was supplemented with 2 mM dibutyryl cyclic AMP (dc-AMP: Santa Cruz, sc-201567A). Media was changed every other day with fresh aliquots of RA, BDNF and dcAMP. Cells were harvested at day 15 for downstream analysis.

**MitoTracker Green FM live imaging**

Light sheet fluorescence microscopy was used to image mitochondrial movement. Cells were incubated with 100 nM of MitoTracker Green FM (Invitrogen, Cat. # M7514) for 20 minutes at 37°C and then washed with prewarmed media prior to imaging. Mitochondrial movements along the neurites were captured every at 0.5 µm z-steps, 10 s intervals, for 3 minutes on a ASI's RAMM frame (ASI), equipped with Mizar TILT light sheet illumination module (Mizar Imaging LLC), Prime 95B sCMOS (Photometrics) camera, and a Plan Apo TIRF 100x/1.45 NA oil (Nikon) objective. Prior to fluorescence imaging, a brightfield image was captured to confirm presence of neurites. Kymographs of mitochondrial movement along neurites were generated using the line tools and resliced command in FIJI.

### Plasmids and siRNAs

Plasmids encoding vinculin tomato or vimentin-GFP (Addgene #56439) were transfected with Lipofectamine 3000 in Opti-Mem (ThermoFisher) according to the manufacturer's instructions. To induce vimentin bundle cells were treated for 4 or 24 hours with 10 µm simvastatin, in their standard culture medium. For *PTEN* knockdown a combination of four siRNAs targeting *PTEN* were used (ON-TARGETplus Human PTEN [5728] siRNA - SMARTpool: Target sequence1#: Sense GAUCAGCAUACACAAAUUA, Antisense: UAAUUUGUGUAUGCUGAUC; Target sequence2#: Sense GACUUAGACUUGACCUAUA, Antisense: UAUAGGUCAAGUCUAAGUC; Target sequence3#: Sense GAUCUUGACCAAUGGCUAA, Antisense: UUAGCCAUUGGUCAAGAUC; Target sequence4#: Sense CGAUAGCAUUUGCAGUAUA, Antisense: UAUACUGCAAAUGCUAUCG). These siRNAs were at a concentration of 10 nm each and were transfected in combination using Lipofectamine 3000 (ThermoFisher), according to the manufacturer's instructions. A negative control siRNA with no significant sequence similarity to human gene sequences was used as a control.

### Fluorescence recovery after photobleaching (FRAP)

Cells were transfected with plasmid encoding vimentin-GFP or vinculin-tomato. 48 hours post transfection FRAP experiments were conducted as described previously using an LSM880 microscope (Zeiss)[6]. 2µm x 2µm regions of interest in cells expressing vimentin-GFP or vinculin-tomato were excited with the 488 nm or 568nm laser lines respectively. Following photobleaching regions of interest were imaged a minimum of for 50 cycles with a 1 second interval. Image and data acquisition were performed using Zen Black and Zen Blue software (Zeiss). Fluorescence intensity in the photo-bleached regions at each time point were quantified as a percentage of fluorescence intensity before photobleaching.

#### Transwell migration assays

Cell migration and invasion abilities were assessed using Transwell cell culture inserts (BD Biosciences). For the cell migration assay, 2.5 $\times 10^4$ cells in 500 µL in serum-free medium were seeded directly into the wells of Transwell chambers with 8 µm-pore membranes. Medium containing 10% FBS, was added into the lower chamber. After 24h cells were fixed and stained with 2% Giemsa blue stain (Sigma). Cells adhering to the upper surface of the membrane were removed using a cotton applicator. Cells on the lower side of the membrane were counted. Five fields were randomly selected per cell line and the mean number of cells quantified.

#### Animal Care

All animal procedures used in this study were approved by the Institutional Animal Care and Use Committee at the University of North Carolina at Chapel Hill. Mice were housed in an AAALAC accredited facility in accordance with the Guide for the Care and Use of Laboratory Animal. All animal procedures were approved by the University of North Carolina at Chapel Hill Institutional Animal Care and Use Committee (IACUC). *SACS*$^{(-/-)}$ mice were a kind gift from Dr. Stefan Strack. *SACS*$^{(+/-)}$ mice were generated by mating *SACS* KO mice with wild type C57BL/6J mice (Jackson Laboratories). Primers for genotyping are as follows: WT allele forward primer: 5' - GCTGTCAGGGGGAAATCTGATAAAG –3', WT allele reverse primer: 5' - GCAGCACCTTTAGACAAAAGATTGC –3', SACS KO allele forward primer: 5' - CAACCTTGGAGAAACTGTGCCTG – 3', SACS KO allele reverse primer: 5' - CACCGACGCCAATCACAAACAC –3'.

#### Tissue collection and immunostaining

Mice were anesthetized using pentobarbital and perfused with 4% PFA in PBS via transcardiac perfusion. Tissue was dissected and drop-fixed in 4% PFA for 24 hours at 4˚C, followed by a 48-hour incubation in X% sucrose at 4˚C. Tissue was mounted in M1 Embedding matrix (ThermoFisher) and stored at -80˚C. Tissue was sectioned on a CryoStar NX50 (ThermoFisher) in 30µm sections. Slices were transferred into a 96 well dish in 100uL of permeabilization buffer (5% NDS, 0.3% Triton X-100, 2% DMSO, 0.01% Sodium Azide, 1X PBS) for 60 minutes on a shaker at room temperature. Following the incubation, buffer was removed and 100 ul of primary antibodies diluted in staining buffer were added for 48 hours. with gentle rocking at room temperature. Sections were washed with PBST (0.1% Triton X-100/1X PBS) three times for 10 minutes, followed by 2 hr. room temperature incubation with gentle rocking in the dark in 100 ul staining buffer with secondary antibodies. The secondary antibody was removed and DAPI diluted to 1:1000 was added for 5 minutes before 3 washes in PBST (0.1% Triton X-100/1X PBS). Slices were mounted on Superfrost Plus slides (ThermoFisher), dried, and ~200ul mounting medium (Sigma, Polyvinyl alcohol mounting medium with DABCO®, antifading) was added prior to placing a cover slip. Slides were dried overnight prior to imaging.

### Sacsin interactome analysis

Proteins were extracted from 8.5x$10^6$ cells of KO and WT SHSY5Y cell lines using 500 µl of the lysis buffer (RIPA with phosphatase and protease inhibitors (EDTA free, Merk). The lysates were cleared by centrifugation (13000g) for 15 minutes at 4°C and then incubated with 1/50 dilution of anti-sacsin antibody (Abcam, ab181190) overnight at 4°C with slow rotation. Protein A Dynabeads (Merk) were equilibrated with the ice-cold lysis buffer and incubated with each cell lysate/antibody mix for 2 hours at 4°C with slow rotation. The beads were then washed with the ice-cold lysis buffer and the bound proteins were eluted with 40µl of 2xLaemmli buffer. The samples were then heated at 95°C for 5 minutes, centrifuged and loaded onto the NuPAGE™ 4 to 12%, Bis-Tris, 1.5 mm, Mini Protein Gel (ThermoFisher). The gels were resolved in 1x MOPC for 2 hours at 4°C and then stained with SimplyBlue SafeStain (ThermoFisher) according to the manufacturers protocol. Each gel lane was then cut into 10 fragments and the proteins were extracted from the gel using trypsin digest protocol as described as previously described[88]. Digests were analysed using a Waters NanoAcquity Ultra-Performance Liquid Chromatography system and data processed using using PLGS v3.0.2 (Waters, UK). For identification of proteins that interact specifically with sacsin, those identified in both KO and WT samples were considered as binding non-specifically and subtracted from the list of putative sacsin interactors.

### Materials and Methods References


83 Arend, K. C. *et al.* Kinome Profiling Identifies Druggable Targets for Novel Human Cytomegalovirus (HCMV) Antivirals. *Mol Cell Proteomics* **16**, S263-S276, doi:10.1074/mcp.M116.065375 (2017).

84 Ritchie, M. E. *et al.* limma powers differential expression analyses for RNA-sequencing and microarray studies. *Nucleic acids research* **43**, e47, doi:10.1093/nar/gkv007 (2015).

85 Metz, K. S. *et al.* Coral: Clear and Customizable Visualization of Human Kinome Data. *Cell Syst* **7**, 347-350 e341, doi:10.1016/j.cels.2018.07.001 (2018).

86 Gene Ontology, C. The Gene Ontology resource: enriching a GOld mine. *Nucleic acids research* **49**, D325-D334, doi:10.1093/nar/gkaa1113 (2021).

87 Kuo, J. C., Han, X., Yates, J. R., 3rd & Waterman, C. M. Isolation of focal adhesion proteins for biochemical and proteomic analysis. *Methods Mol Biol* **757**, 297-323, doi:10.1007/978-1-61779-166-6_19 (2012).

88 Patel, V. J. *et al.* A comparison of labeling and label-free mass spectrometry-based proteomics approaches. *Journal of proteome research* **8**, 3752-3759, doi:10.1021/pr900080y (2009).
